# A broad response to intracellular long-chain polyphosphate in human cells

**DOI:** 10.1101/2020.05.01.056192

**Authors:** Emma Bondy-Chorney, Iryna Abramchuk, Rawan Nasser, Charlotte Holinier, Alix Denoncourt, Kanchi Baijal, Liam McCarthy, Mireille Khacho, Mathieu Lavallée-Adam, Michael Downey

## Abstract

Polyphosphates (PolyP) are composed of long chains of inorganic phosphates linked together by phosphoanhydride bonds. They are found in all kingdoms of life, playing roles in cell growth, infection, and blood coagulation. A resurgence in interest in polyP has shown links to diverse aspects of human disease. However, unlike in bacteria and lower eukaryotes, the mammalian enzymes responsible for polyP metabolism are not known. Many studies have resorted to adding polyP to cell culture media, but it is not clear if externally applied polyP enters the cell to impact signaling events or whether their effect is mediated exclusively by extracellular receptors. For the first time, we use RNA-seq and mass spectrometry to define a broad impact of polyP produced inside of mammalian cells via ectopic expression of the *E. coli* polyP synthetase Ppk1. RNA-seq demonstrates that Ppk1 expression impacts expression of over 350 genes enriched for processes related to transcription and cell motility. Analysis of proteins via label-free mass spectrometry identified over 100 changes with functional enrichment in cell migration. Follow up work suggests a role for internally-synthesized polyP in promoting activation of mTOR and ERK1/2-EGR1 signaling pathways implicated in cell growth and stress. Finally, fractionation analysis shows that polyP accumulated in multiple cellular compartments and was associated with the relocalization several nuclear/cytoskeleton proteins, including chromatin bound proteins DEK, TAF10, GTF2I and translation initiation factor eIF5b. Our work is the first to demonstrate that internally produced polyP can activate diverse signaling pathways in human cells.

**Significance Statement:** For many years following its discovery in 1890, polyphosphates (polyP) were dismissed as evolutionary fossils. Best understood for its role in bacteria and yeast, our understanding of polyP in mammals remains rudimentary because the enzymes that synthesize and degrade polyP in mammalian systems are currently unknown. In our work, we carried out large-scale transcriptome and proteome approaches on human cells designed to accumulate internally produced polyP via ectopic expression of a bacterial polyP synthetase. Our work is the first to systematically assess the impact of increased intracellular polyP.

## Introduction

Polyphosphates (polyP) are molecules comprised of inorganic phosphates joined by high energy phosphoanhydride bonds in chains of 3-1000 units in length. Although polyP itself appears to be present across all kingdoms of life, its mechanisms of synthesis appear poorly conserved. In bacteria, polyP is synthesized by the Ppk1 enzyme that uses ATP as a substrate (Ahn and Kornberg, 1990). In the enterobacterium *E. coli* in particular, polyP levels are low during logarithmic growth in rich media but increase dramatically to approximately 20 mM (measured in terms of individual phosphate units) in response to a variety of stresses such as amino acid starvation (Rao et al., 1998). These chains range from 100-1000 units (Kornberg et al., 1999; Kumble et al., 1996). PolyP accumulation is countered by Ppx, an exopolyphosphatase that degrades polyP beginning at the end of the chain (Akiyama et al., 1993; Rashid et al., 2000). In yeast, polyP is synthesized by the VTC complex (Hothorn et al., 2009). The VTC complex is situated within the vacuolar membrane and, like Ppk1, synthesizes polyP using ATP as a substrate. PolyP is translocated into the vacuole as it is synthesized (Gerasimaitė et al., 2014), where it accumulates to remarkably high levels (>200 mM total cellular concentration, comprising up to 10% of the dry weight of the cell) (Auesukaree et al., 2004; Kornberg et al., 1999). Lower levels of polyP can be found in other subcellular compartments including the cytoplasm, nucleus, plasma membrane and mitochondria (Gerasimaitė and Mayer, 2016; Lichko et al., 2006). How polyP localizes to these areas of the cell is not understood.

In stark contrast to bacteria and yeast, the enzymes that synthesize polyphosphate in mammalian cells have remained elusive (Desfougères et al., 2020). There are no clear Ppk1 or VTC homologs in mammalian cells. On the other hand, Prune, previously characterized as an interactor of GSK-3β and a regulator of Wnt signaling can degrade polyP *in vitro* (Tammenkoski et al., 2008), but it is not clear if this enzyme has any role in polyP metabolism *in vivo.* The same is true for alkaline phosphatase and three members of the Nudix family of hydrolases, DIPP1-DIPP3, which are homologs of yeast Ddp1 (Lonetti et al., 2011). Whole-cell levels of polyP (25-120 μM – Pi residues) in human cells and tissues (Kumble and Kornberg, 1995) are considerably lower than those measured in bacteria or yeast. Higher local concentrations may exist in certain subcellular compartments. For example, Jimenez-Nunez *et al.* reported high local concentrations of polyP in the nucleus of myeloma cells (Jimenez-Nuñez et al., 2012). PolyP also accumulates to levels of 130 mM in the dense granules of platelets (Ruiz et al., 2004), a lysosome-related organelle similar in many ways to the yeast vacuole. Release of polyP from these structures is thought to play an important role in the activation of the blood coagulation cascade (Travers et al., 2015). The accumulation of polyP in these structures may suggest elements of conservation with yeast in terms of polyP synthesis or storage.

With no enzymes responsible for *in vivo* polyP synthesis and degradation identified, studying polyP in mammalian systems presents a unique challenge. Several groups have expressed yeast Ppx1 exopolyphosphatase enzyme in mammalian cells in an attempt to deplete endogenous polyP. Wang *et al.* first used this approach to study the role of polyP role in mammalian target of rapamycin (mTor) activation in MCF-7 mammary cancer cells (Wang et al., 2003). Subsequently others have used ectopic expression of Ppx1 to study the role of PolyP in mitochondria (Abramov et al., 2007; Seidlmayer et al., 2012), wound healing (Simbulan-Rosenthal et al., 2015), the DNA damage response (Bru et al., 2017), and glycolysis (Nakamura et al., 2018).

The impact of increased polyP has been studied by adding polyP chains of various lengths directly to culture media (Dinarvand et al., 2014; Hassanian et al., 2015; Holmström et al., 2013; Xie et al., 2019). For example, previous work from the Rezaie group showed that polyP chains of 70 units interact with the RAGE and P2Y1 receptors on the plasma membrane and activates the mTORC1 and TORC2 pathways in endothelial cells (Dinarvand et al., 2014; Hassanian et al., 2015). In these and other studies it is not clear whether polyP is entering the cell and/or whether this is required for the observed effects. Indeed, the requirement of external receptors for activation of these pathways suggests that polyP may be acting extracellularly (Dinarvand et al., 2014; Hassanian et al., 2015). In this study, we investigate the impact of internally-synthesized polyP in mammalian cells produced via ectopic expression of the *E. coli ppk1+* gene. We report that ectopic expression of the Ppk1 protein results in polyP accumulation throughout the cell, including the nucleus. Using transcriptomic and proteomic analyses, we demonstrate a broad impact of polyP on diverse pathways, including activation of the ERK1/2-EGR1 signaling axis. PolyP accumulation also results in redistribution of chromatin bound proteins DEK, TAF10 and GTF2I, and translation initiation factor eIF5b from the nucleus/cytoskeleton to the cytoplasm. Our work will serve as a novel resource to interrogate polyP biology in higher eukaryotes.

## RESULTS

### Internal synthesis of polyP in mammalian cells

To determine the impact of polyP accumulation in mammalian cells, we transiently transfected either an empty vector (control) or a vector expressing the *E. coli ppk1+* gene under the CMV promoter **(Fig. 1A)**. The *ppk1+* gene encodes Ppk1, the enzyme responsible for synthesizing polyP chains in *E. coli* (Ahn and Kornberg, 1990; Kornberg et al., 1956). Ppk1 expression in mammalian cells has been used previously as a gene reporter system in magnetic resonance imaging techniques (Ki et al., 2007). We also previously used this experimental set-up to identify the first human targets of polyphosphorylation, a recently described post-translational modification wherein polyP chains are covalently attached to lysine residues (Bentley-DeSousa et al., 2018). However, we did not directly examine polyP production or downstream impacts of its accumulation. Ppk1 expression resulted in accumulation of long-chain polyP (>130 residues) in HEK293T cells **(Fig. 1B, lanes 6 and 7)**. The size distribution of polyP produced is similar to the polyP made in *E. coli* under starvation (stress), and contrasts with the shorter and more heterogeneous chains synthesized by the VTC complex in yeast **(Fig. 1B, lanes 8 to 11)**. As we demonstrated previously, Ppk1 expression caused a dramatic decrease in electrophoretic mobility for polyphosphorylation targets Nucleolin and MESD when analyzed on NuPAGE gels **(Fig. 1C)**. This change in electrophoretic mobility, visible on NuPAGE, but not SDS-PAGE gels, is a hallmark of lysine polyphosphorylation (Azevedo et al., 2015; Bentley-DeSousa et al., 2018).

**Figure 1.**
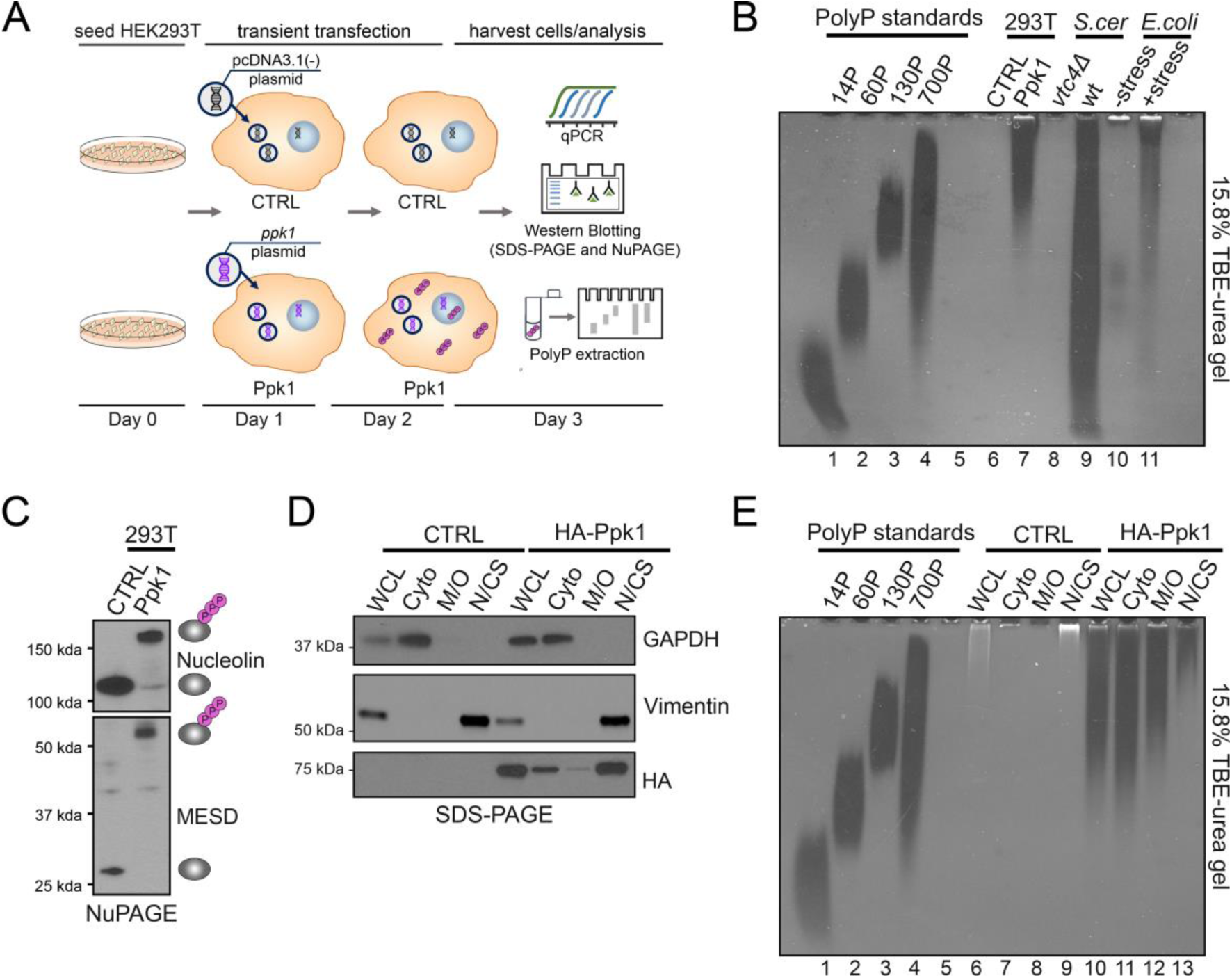
Characteristics of cells that accumulate internal polyP. *(A)* HEK293T cells were seeded on day zero and on day 1 were transfected with either empty vector (pcDNA3.1(-)) or *ppk1+* expression plasmid (pcDNA3.1(-)*ppk1+*) to generate control or Ppk1 conditions, respectively. After 48 hours cell lysates were harvested and analyzed with follow up experiments including, but not limited to qPCR, western blotting (SDS-PAGE and NuPAGE), and polyP extraction. *(B)* PolyP extractions analyzed on a 15.8% TBE-urea gel stained with DAPI from control and Ppk1-expressing HEK293T cells, wt and *vtc4Δ* yeast, and *E. coli* in unstressed or stressed conditions (amino acid starvation via growth in MOPS minimal media). PolyP standards [14P, 60P, 130P (Regentiss, Japan) and 700P (Kerafast)] are presented for comparison. Image shows results from one biological replicate, which is representative of N=3. *(C)* NuPAGE analysis of known polyphosphorylated proteins. Total protein was extracted using RIPA buffer prior to NuPAGE analysis via western blotting (same blot) with indicated antibodies. *(D)* 48 hours after transfection HEK293T cells were fractionated using Cell Signaling Technologies Cell Fractionation Kit according to manufactures protocol. SDS-PAGE analysis (4–20% Criterion TGX Stain-Free Protein Gel) of whole cell lysate (WCL) and cell fractions from control and Ppk1 conditions with antibodies known to be localized to specific fractions: Cytoplasm (Cyto) – GAPDH and Nuclear/Cytoskeleton (N/CS) – Vimentin. Image is representative of N≤3. Representative image shown here is from the same biological sample from different blots analyzed in parallel. *(E)* PolyP extractions analyzed on a 15.8% TBE-urea gel from cell fractionations of control and Ppk1 conditions HEK293T cells. PolyP was extracted directly from cell fraction aliquots either immediately following fractionation or from fractions stored at −80 °C.

Next, we investigated the intracellular localization of the recombinant Ppk1 protein. As there are no commercially available antibodies for *E. coli* Ppk1 protein, we expressed an HA-tagged Ppk1 and carried out cell fractionation experiments. Commonly used cell fractionation markers showed expected separation for control and HA-Ppk1 samples **(Fig. 1D)**. We found that some HA-Ppk1 protein was localized in each of these compartments, with the majority observed in the cytoplasmic and nuclear/cytoskeletal fractions **(Fig. 1D).** Surprisingly, polyP concentration did not correlate with the level of HA-Ppk1 expression. While polyP was observed in all fractions, the highest concentrations were in the cytoplasm and the lowest in the nuclear/cytoskeletal fractions **(Fig. 1E, lanes 10 - 13)**. Nevertheless, these results demonstrate that mammalian cells are capable of supporting the ectopic expression of Ppk1 and the accumulation of polyP *in vivo*.

### No significant impact of polyP on cell-cycle, apoptosis or autophagy markers

With a system in place to successfully induce accumulation of polyP inside mammalian cells, we set out to determine the impact of elevated and internally-synthesized polyP on common cell signaling pathways. PolyP is important for cell cycle regulation in yeast and bacteria (Bru et al., 2016; Racki et al., 2017). Coordination of the cell cycle is a highly complex process involving the activities and levels of numerous proteins, including cyclins (Ingham and Schwartz, 2017; Schafer, 1998). Altered levels of cyclins serve as an indicator of cell-cycle changes (Malumbres and Barbacid, 2005; Yam et al., 2002). However, western blotting analysis revealed no apparent changes in cyclins protein levels when Ppk1 was expressed as compared to control, suggesting no gross disruption of normal cell cycle activities **(Supplemental Fig. 1A and B)**. Next, we examined common markers of apoptosis (programmed cell death), which is regulated by a balance between pro- and anti-apoptotic pathways, including transcriptional regulation (Kumar and Cakouros, 2004; Portt et al., 2011). Indeed the external application of polyP has been reported to induce apoptosis in select human plasma cells (Hernandez-Ruiz et al., 2006) or under specific conditions, such as in combination with chemotherapeutic agents (Xie et al., 2019). We used RT-qPCR to determine the levels of *p53*, *BAX*, and *CASPASE-9*, three mRNAs commonly involved in the regulation of apoptotic pathways (Cummings, 1996; Liu and Lobie, 2007; Raman et al., 2000). Analysis of this data suggested no consistent change of these three apoptosis-related mRNAs (**Supplemental Fig. 1C)**. Lastly, we tested if we could detect changes in autophagy, a highly-conserved intracellular degradation and recycling system (Shibutani et al., 2015), in Ppk1-expressing cells. Measuring the conversion of LC3-I to LC3-II by immunoblotting is a commonly used method for monitoring autophagy (Mizushima et al., 2010). We saw no significant change in the LC3-I/LC3-II ratio between control and Ppk1-expressing cells **(Supplemental Fig. 1D)**. Taken together, these data suggest that the expression of Ppk1 in HEK293T cells does not significantly disrupt cell cycle progression and does not induce apoptosis, or autophagy. To take a broad, unbiased look at the impact of Ppk1 expression, we used RNA-sequencing (RNA-seq) and label-free mass spectrometry to uncover changes in gene and protein expression.

### Hundreds of differentially expressed genes in cells making excess polyP

To investigate the impact of polyP accumulation on the transcriptome, we carried out a large-scale RNA-seq analysis in HEK293T cells **(Fig. 2A)**. NuPAGE analysis of known polyphosphorylated proteins DEK and MESD were used to confirm polyP production in Ppk1 samples used for RNA-seq **(Fig. 2B)**. We found 313 downregulated and 47 upregulated genes (FDR-adjusted p-value<0.05, log_2_-fold change threshold > 1.25 and < 0.8) in Ppk1-expressing cells **(Fig. 2C and Supplemental Table 2)**. Among the top protein-coding genes that showed significant downregulation in Ppk1-expressing cells were three members of the Inhibitor of DNA-binding/differentiation (ID) family (*ID1*, *ID2*, *ID3*), *CCR4*, *DHRS2*, and *SFRP4*. Included among the genes that showed significant upregulation included *MYH15*, *VGF*, *ETV5*, and multiple members within the same protein family, such as the Early Growth Response (EGR) transcription factor family (*EGR1*, *EGR3*). Notably, top-ranked down and upregulated genes also included pseudogenes and lncRNAs **(Supplemental Table 2)**. To validate hits from our RNA-seq we carried out RT-qPCR on 12 candidate targets that were predicted to have differential expression based on the RNA-seq data **(Fig. 2D - F and Supplemental Table 2 and 3)**. All results shown are normalized to controls within paired biological replicates. In total, 9 out of the 12 genes tested showed statistically significant differences in the direction that was predicted by the RNA-seq data including *HIST1H1D*, *ETV5*, *ID3*, and *CCR4*, resulting in a validation rate of 75% **(Fig. 2D - F)**. Additionally, we confirmed that 2 genes that we worked with previously, *NCL* and *NOP56* (Bentley-DeSousa et al., 2018), did not show any significant differences in differential expression, as predicted by the RNA-seq analysis **(Fig. G)**. We conclude that our dataset is of high quality and that polyP production in mammalian cells stimulates significant reprograming of gene expression. GO-term enrichment analysis performed on all significantly differentially expressed genes revealed 11 enriched terms **(Fig. 2H and Supplementary Table 2**; FDR-adjusted p-value < 0.05). The top 4 most significantly enriched GO-terms were associated with DNA, specifically transcription regulation and DNA binding. Altogether, these results show significant changes in the transcriptome in cells accumulating internally-synthesized polyP.

**Figure 2.**
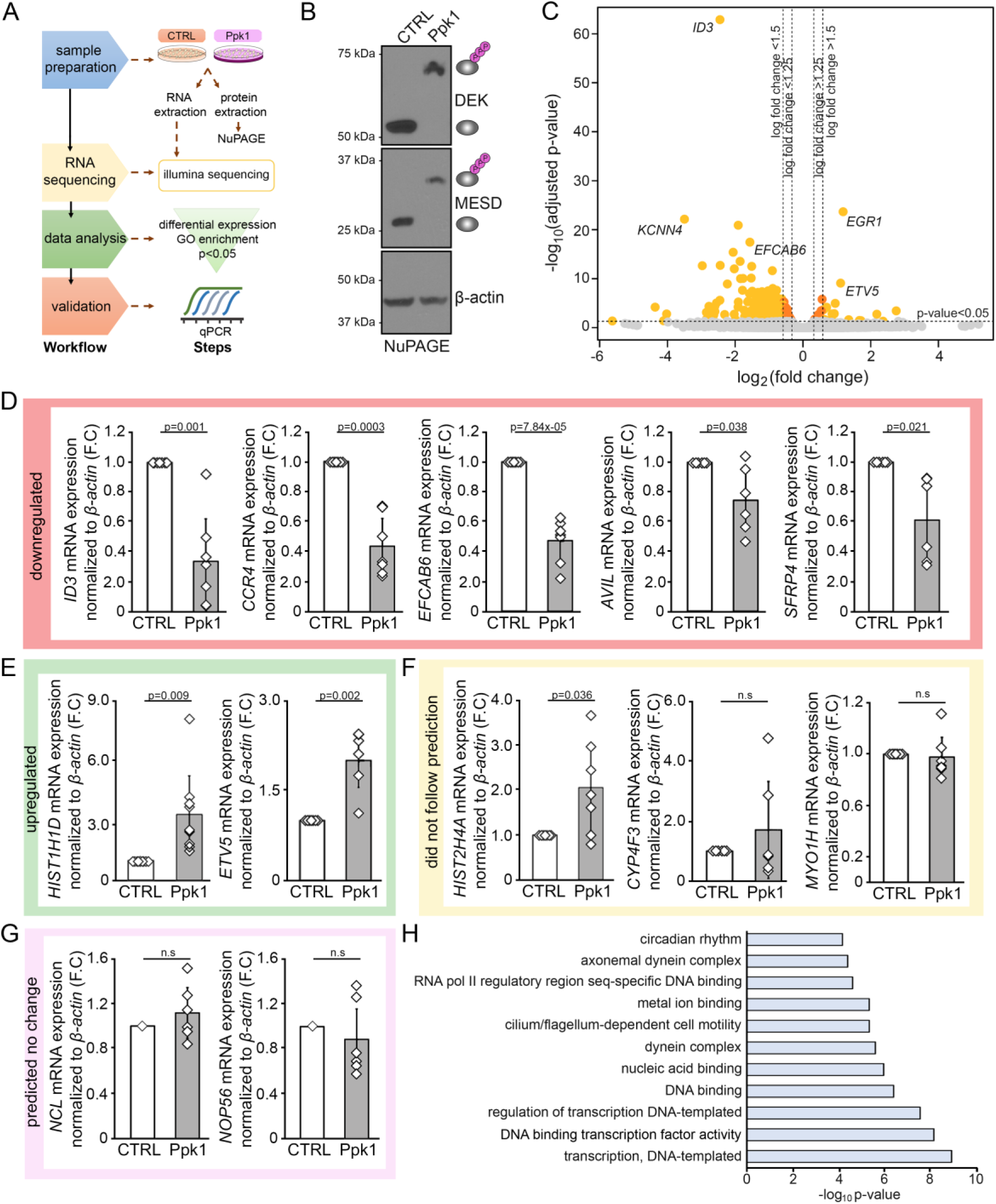
RNA-seq analysis of cells that accumulate polyP. *(A)* Workflow of RNA-seq analysis. 48 hours after transfection with either control or *ppk1+* expression plasmid, RNA and protein were harvested from cells. RNA was immediately shipped to Genome Quebec for RNA-seq. RNA-seq data was analyzed and a small subset of genes were validated by RT-qPCR (details in materials and methods). *(B)* NuPAGE analysis showing polyphosphorylation induced shifts of DEK and MESD in lysates harvested from cells sent for RNA-seq. β-actin, a non-polyphosphorylated protein, was used as a loading control. Image is representative of N=3. Representative image shown here is from the same biological sample from different blots analyzed in parallel (MESD and β-actin were analyzed on one blot separate from DEK). *(C)* Volcano plot shows log_2_(fold change) and log_10_(FDR-adjusted p-value) for all genes obtained from the RNA-seq experiment. Genes were classified as significant if their FDR-adjusted p-value<0.05. Statistical analysis of RNA-seq data is included in detail in the materials and methods. *(D,E,F,G)* RT-qPCR validation of top hits from RNA-seq differential expression analysis. Changes in mRNA levels are represented by fold change (F.C). *(D,E)* Hits that were found to show expression changes in the same direction as predicted by the RNA-seq data are shown inside green (up) or pink (down) boxes. *(F)* Hits that did not show expression changes that followed predicted patterns from RNA-seq data are inside a yellow box. *(G)* Graphs inside the light pink box are hits that were predicted and found by RT-qPCR not to show significant expression changes from control to Ppk1 conditions. Primers used for RNA-seq validation are included in **Supplemental Table 3**. Graphs represent N≤4 for each mRNA. P-values are shown as numerical values or if p≤0.05 as non-significant (n.s). Statistical tests performed were one-sample t-Test (unequal variances) where error bars represent standard deviation. *(H)* GO-term analysis of differentially expressed genes (FDR-adjusted *p*-value<0.05). GO-term analysis is described in materials and methods.

### Global changes in protein levels in polyP producing cells

We next investigated if changes in gene expression were accompanied by changes in protein levels. We used label-free mass spectrometry to measure global protein levels in five biological replicates of HEK293T cells transiently transfected with either Ppk1 or control plasmids **(Fig. 3A)**. NuPAGE analysis was again used to test for polyphosphorylation of DEK and MESD to confirm production of polyP in samples processed for mass spectrometry **(Fig. 3B)**. Our analysis of the mass spectrometry data revealed statistically significant changes in the levels of 14 proteins in cells with Ppk1 expression as compared to control **(Fig. 3C; FDR<15%)**. Importantly, we detected our recombinant Ppk1 protein as present only in Ppk1 conditions **(Supplemental Table 4)**. Amongst the proteins identified was CUL1, two Minichromosome Maintenance Complex Component (MCM) proteins MCM3 and MCM4, and RRBP1. We further characterized an additional group we called “singletons”, composed of proteins that show changes in their levels but could not be included in our statistical analysis as they were not detected in all five biological replicates in both conditions **(Fig. 3D, details defining singletons in materials and methods)**. This group included Dehydrogenase/Reductase 2 (DHRS2), a protein encoded by the *DHRS2* gene, which was also one of the hits we uncovered in our RNA-seq to be decreased with Ppk1 expression **(Supplemental Table 2)** and discussed further below. Further comparative analysis revealed 95.6% of the proteins detected in our mass spectrometry did indeed have their corresponding RNAs detected in our RNA-seq. Interestingly, DHRS2 and DHCR24 were the only hits from the mass spectrometry that also showed significant changes in the RNA-seq. This data suggests that the proteomic changes we observe with Ppk1 expression may be occurring at a post-transcriptional level. GO-term enrichment analysis of the significantly differentially expressed and singletons proteins revealed only one significantly enriched term, positive regulation of epithelial cell migration (adjusted p-value = 0.028).

**Figure 3.**
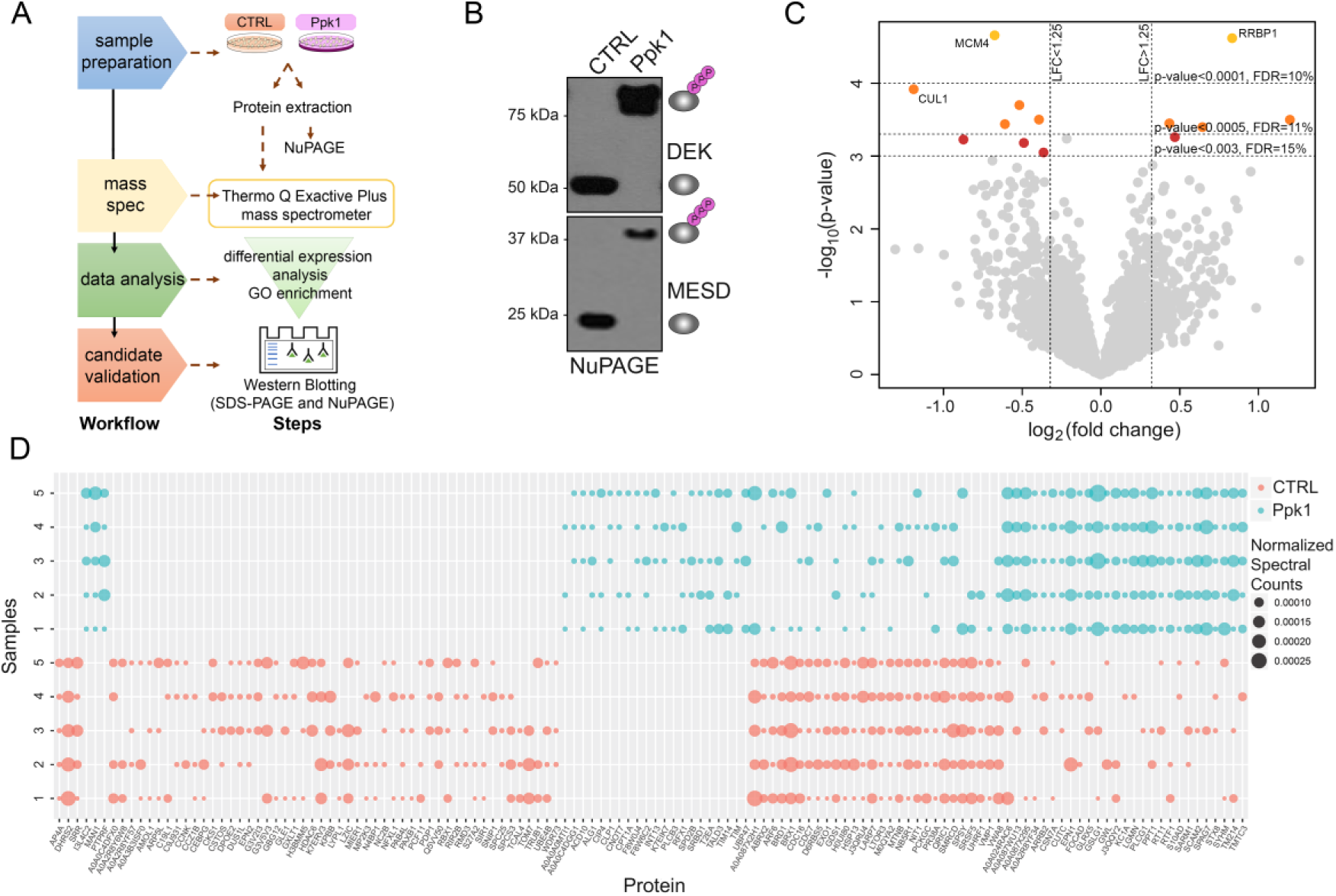
Global changes in protein levels in cells that accumulate polyP. *(A)* Workflow of mass spectrometry analysis. 48 hours after transfection with either control or *ppk1+* expression plasmid, protein was harvested from cells and a small portion of sample was used to confirm polyphosphorylation of targets by NuPAGE in all samples. The remaining portion of each sample was immediate shipped to UC Davis for mass spectrometry (details included in materials and methods). *(B)* NuPAGE analysis showing polyphosphorylation induced shifts of DEK and MESD in lysates harvested from cells sent for MS. Image is representative of N=5. Representative image shown here is from the same biological sample from different blots analyzed in parallel. *(C)* Volcano plot shows log_2_(fold change) and -log_10_(p-value) for all proteins identified in all 5 replicates of each condition. Statistical analysis of mass spectrometry data is included in detail in the materials and methods. *(D)* “Singletons” from mass spectrometry data. Categories shown in both control (pink) and Ppk1 (blue) conditions include “All or None” (proteins identified in 5/5 in one condition and 0/5 in the other), 2) “Mostly All or None” (proteins identified in (3 or 4)/5 in one condition and 0/5 in the other), and 3) “All or Mostly None” (proteins identified in 5/5 in one condition and (1 or 2)/5 in the other). Size of bubbles reflect normalized spectral counts, a proxy of the relative abundance of the protein within each biological replicate (samples 1-5), for each condition.

Extracting the protein-protein interactions between the significantly differentially expressed proteins and singletons using STRING (Szklarczyk et al., 2019) revealed several subsets of conserved interaction networks **(Supplemental Fig. 2)**. We further analyzed this STRING network by identifying GO-terms that were associated with the individual nodes with this network. From this, we saw several dense subnetworks where the majority of proteins within the nodes were associated with a specific GO-term, including RNA processing, DNA replication, and transcription elongation from the RNA polymerase II promoter. In these cases, very few proteins associated with these terms were identified outside of the isolated nodes. Finally, we also saw that a number of the proteins within our overall STRING-derived network were associated with the GO-terms nucleic acid metabolic processes and cellular nitrogen compound metabolic processes **(Supplemental Fig. 2)**. Altogether, these results confirm that internal polyP production is associated with changes at the proteome level.

### Internal synthesis of polyP leads to downregulation of the dehydrogenase/reductase DHRS2

Taken together, our RNA-seq and mass spectrometry data reveals global changes in both the transcription and proteome of Ppk1-expressing cells. To extend and further validate these analyses, we followed up on several of these targets including one that came out as a top hit in both our RNA-seq and mass spectrometry experiments, *DHRS2*. *DHRS2* is a gene that encodes for the DHRS2 protein, also known as Hep27 (Donadel et al., 1991; Gabrielli et al., 1995). DHRS2 is a member of the short-chain dehydrogenases/reductases family and has been reported to catalyze the NADPH-dependent reduction of specific dicarbonyl compounds (Shafqat et al., 2006). Data from our RNA-seq revealed that *DHRS2* mRNA levels were decreased by 3.44 fold (deseq. adjusted p-value = 2.90 × 10-14) in Ppk1-expressing cells **(Supplemental Table 2)**. Validation by RT-qPCR supported this expression level change, as *DHRS2* was downregulated by 3.47 fold (p-value = 0.013) with Ppk1 expression **(Fig. 4A)**. To investigate whether the decrease in the *DHRS2* mRNA levels was also reflected in the protein levels, we examined DHRS2 protein changes by western blot **(Fig. 4B)**. In confirmation of our mass spectrometry data **(Fig. 3D and Supplemental Table 4)** we observed a significant decrease in DHRS2 protein, as compared to normalized control **(Fig. 4B and C)**. DHRS2 is reported to stabilize p53 levels via Ser15 phosphorylation in esophageal squamous-cell carcinoma cells (Zhou et al., 2018). However, we observed no changes in p53 Ser15 phosphorylation in Ppk1-expressing cells, as compared to controls **(Fig. 4B and D)**. This difference in our results and previously published data may stem from the use of different cell lines, as detailed in the discussion. It would be interesting to test the impact of polyP-induced DHRS2 downregulation on additional pathways regulated by DHRS2, such as dicarbonyl reduction (Shafqat et al., 2006).

**Figure 4.**
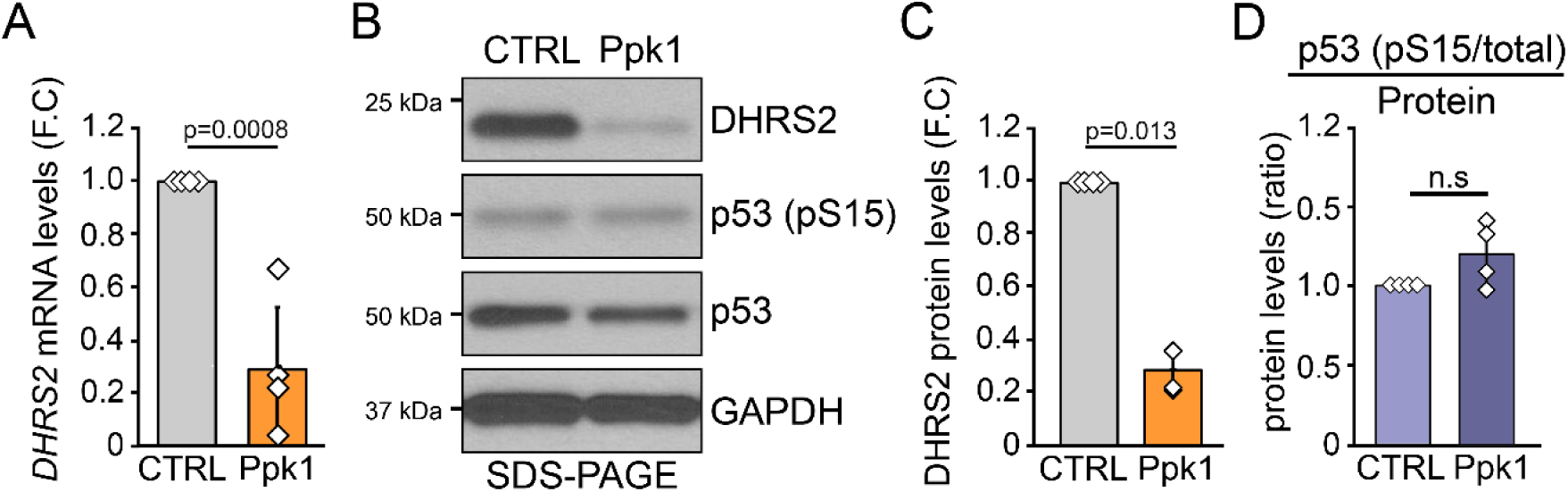
Accumulation of polyP leads to upregulation of DHRS2. RNA and protein were harvested from HEK293T cells following a 48-hour transient transfection of either a control or *ppk1+* expression plasmid. *(A)* RT-qPCR analysis of *DHRS2* mRNA in control and Ppk1 conditions. Changes in mRNA levels are represented by fold change (F.C). *(B)* Western blot analysis with SDS-PAGE (4–20% Criterion TGX Stain-Free Protein Gel) of proteins in control and Ppk1 conditions using antibodies against DHRS2, p53, and phosphorylated p53 (Ser-15). GAPDH was used as a loading control. Image is representative of N≤3. Representative image shown here is from the same biological sample from different blots analyzed in parallel. *(C,D)* Semi-quantitative analysis of protein levels shown in panel B. P-values are shown as numerical values or if p≤0.05 as non-significant (n.s). Statistical tests performed were one-sample t-Test (unequal variances) where error bars represent standard deviation. All experiments shown here are representative of N≤3.

### PolyP synthesis activates ERK1/2 signaling

Analysis of our RNA-seq data revealed that upon Ppk1 expression, *EGR1* levels were increased by 2.28 fold (deseq. adjusted p-value = 2.10-24) **(Supplemental Table 2)**. EGR1 is a transcription factor that plays roles in many cellular pathways including cell growth, differentiation, and survival (Adamson and Mercola, 2002; Milbrandt, 1987; Sukhatme et al., 1988; Veyrac et al., 2014). Analysis by RT-qPCR and western blotting demonstrated that both EGR1 mRNA and protein levels were increased with Ppk1 expression **(Fig. 5A - C)**. *EGR1* mRNA transcription is regulated by activation of the ERK1/2 pathway (Roskoski Jr, 2012). Indeed, we found that Ppk1 expression induced ERK1/2 phosphorylation on tyrosine residues Y202 and Y204, without impacting overall ERK1/2 levels **(Fig 5. D and E)**. These residues are phosphorylated by the MEK1/2 kinases in the signaling cascade that depends on upstream Raf and Ras signaling. Interestingly, Hassanian et al. previously reported that incubation of endothelial cells with external polyP induced the phosphorylation of ERK1/2 (Hassanian et al., 2015). They also found that this same treatment resulted in activation of the mTORC1 pathway as measured by phosphorylation of p70S6K (Hassanian et al., 2015). Likewise, we found that p70S6K was phosphorylated in cells expressing Ppk1 **(Fig. 5 F and G)**. For the first time, our work demonstrates that internal production of polyP can activate conserved pathways that impinge on cell growth and development.

**Figure 5.**
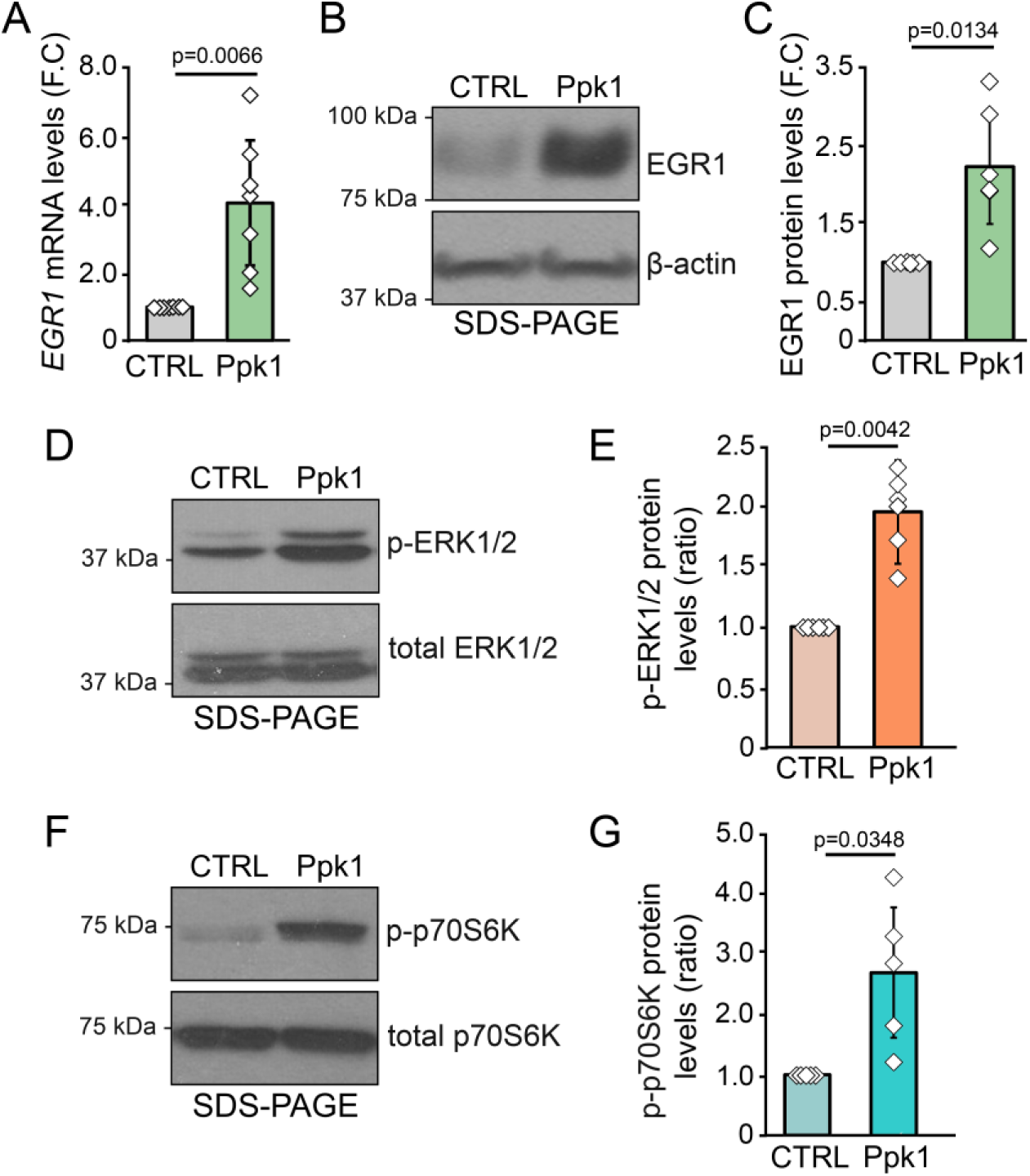
Activation of ERK1/2 signaling by production of polyP. RNA and protein were harvested from HEK293T cells following a 48-hour transient transfection of either a control or *ppk1+* expression plasmid. *(A)* RT-qPCR of *EGR1* mRNA levels in control and Ppk1 conditions. Changes in mRNA levels are represented by fold change (F.C). *(B)* western blotting analysis by SDS-PAGE (4–20% Criterion TGX Stain-Free Protein Gel) in control and Ppk1 conditions cells using the EGR1 antibody with β-actin as a loading control. Image is representative of N≤3. Representative image shown here is from the same biological sample from different blots analyzed in parallel. *(C)* Semi-quantitative analysis of protein levels shown in panel B. *(D)* Western blotting analysis by SDS-PAGE in control and Ppk1 conditions cells using antibodies against total ERK1/2 and phosphorylated ERK1/2. Image is representative of N≤3. Representative image shown here is from the same biological sample from different blots analyzed in parallel. *(E)* Semi-quantitative analysis of protein levels shown in panel D. *(F)* Western blotting analysis by SDS-PAGE in control and Ppk1 conditions cells using antibodies against total p70S6K and phosphorylated p70SK6. Image is representative of N≤3. Representative image shown here is from the same biological sample from different blots analyzed in parallel. *(G)* Semi-quantitative analysis of protein levels shown in panel G. P-values are shown as numerical values or if p≤0.05 as non-significant (n.s). Statistical tests performed were one-sample t-Test (unequal variances), where error bars represent standard deviation.

### Relocalization of nuclear proteins in cells accumulating polyP

Changes in protein localization can dictate a protein’s function and/or activity independently of absolute protein levels (Bauer et al., 2015). Changes in protein localization are often dictated by post-translational modifications. Indeed, work in yeast has demonstrated that polyphosphorylation of RNA binding protein Nsr1, and DNA topoisomerase Top1 modulates the distribution of both proteins between the nucleoplasm and the nucleolus (Azevedo et al., 2015). To examine the impact of polyP accumulation in HEK293T cells on protein localization, we tested whether Ppk1 expression altered the localization of the known polyphosphorylated proteins NOP56, eIF5b, GTF2I, and DEK (Azevedo et al., 2018; Bentley-DeSousa et al., 2018). Upon HA-Ppk1 expression, NOP56 did not show any localization changes from control to HA-Ppk1 conditions **(Fig. 6A)**. Interestingly, eIF5b and GTF2I both showed decreases in protein amount present in the nuclear/cytoskeletal fraction when Ppk1 was expressed **(Fig. 6A and B)**. Fractionation markers for the representative biological replicate in **Fig. 6B** are shown in **Fig. 1D**. DEK, in agreement with previous studies (Hu et al., 2007; Kappes et al., 2001), was localized to the nuclear/cytoskeletal fraction in control conditions. In contrast, in HA-Ppk1-expressing cells, DEK showed a dramatic redistribution to other cellular compartments **(Fig. 6B)**. Based on these findings, we expanded our results to include a screen of 11 additional proteins, using available antibodies targeting both polyphosphorylated and non-polyphosphorylated targets **(Fig. 6 and Materials and Methods)**. Most proteins showed identical fractionation patterns in control and Ppk1-expressing cells. This result suggests that re-distribution of proteins in cells making excess polyP is specific to individual proteins rather than a general phenomenon. However, we did find a consistent decrease in the nuclear/cytoskeletal fraction of one non-polyphosphorylation target, TAF10, following Ppk1 expression **(Fig. 6A)**. PolyP concentrations in all compartments were high enough to confer polyphosphorylation of DEK **(Fig. 6C)**. Taken together, these results show that internal accumulation of polyP alters protein localization of both targets and non-targets of polyphosphorylation in mammalian cells.

**Figure 6.**
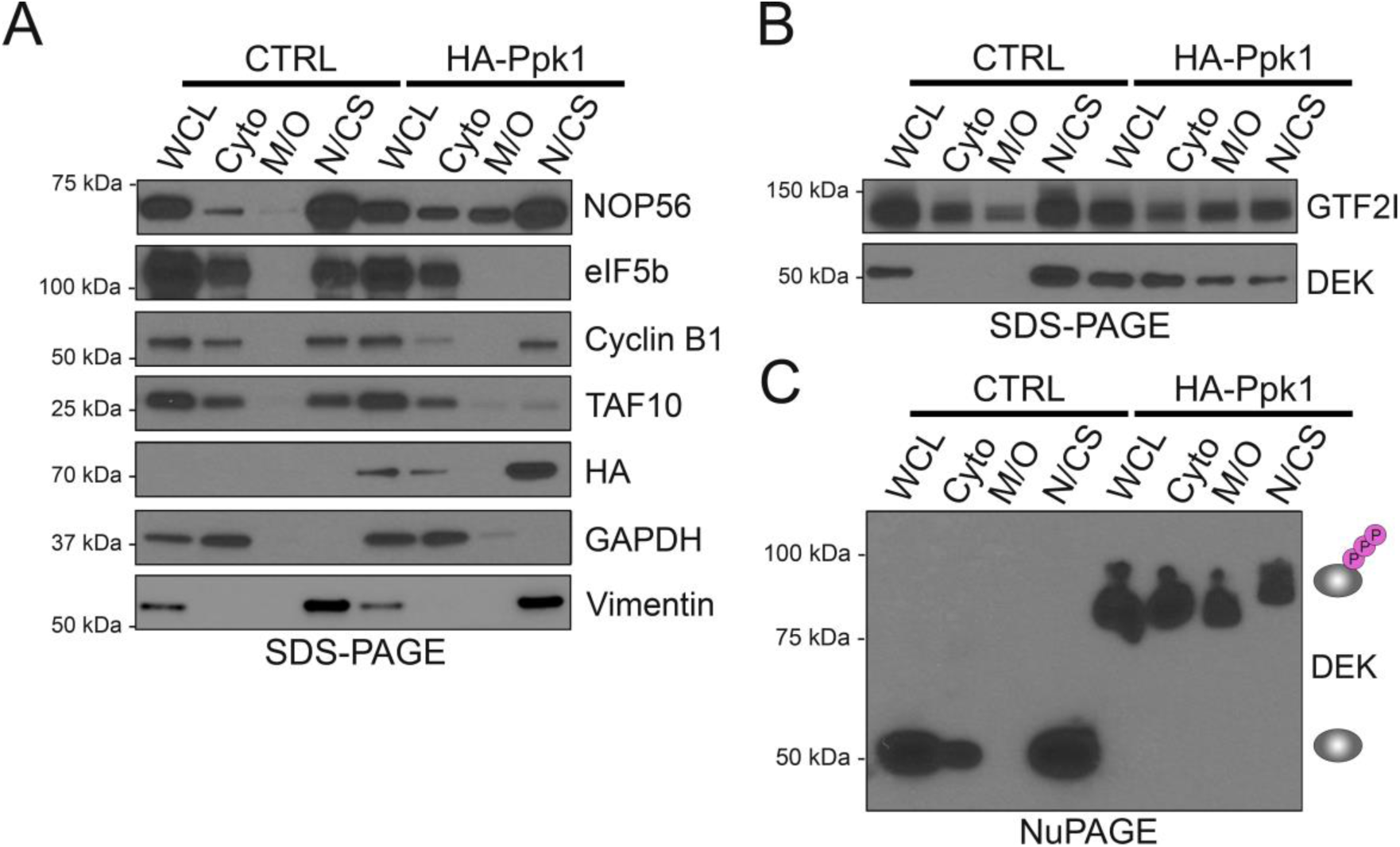
Changes in protein localization in both polyphosphorylated and non-polyphosphorylated proteins with polyP accumulation. HEK293T cells were transiently transfected for 48 hours with either a control or *ppk1+* expression plasmid. *(A, B)* Fractionation of cells was performed using Cell Signaling Technologies Cell Fractionation Kit according to manufactures protocol. SDS-PAGE analysis (4–20% Criterion TGX Stain-Free Protein Gel) of whole cell lysate (WCL) and cell fractions from control and Ppk1 conditions with antibodies against known polyphosphorylated proteins NOP56, eIF5b, GTF2I, and DEK and non-polyphosphorylated proteins, Cyclin B1 and TAF10 were also assessed using antibodies against indicated proteins. Image is representative of N≤3. Representative image shown here is from the same biological sample from different blots analyzed in parallel. Amount of cell material used to generate each fraction was consistent between all biological replicates tested. Fractionation markers for panel *B* are shown in Fig. 1D. *(C)* NuPAGE analysis of cell fractionations from HEK293T control and Ppk1 conditions of the polyphosphorylated protein target DEK. Image is representative of N=3.

## Discussion

PolyP is a molecule that plays many roles across all kingdoms of life (Kulaev and Vagabov, 1983). PolyP research in mammalian cells is hampered by a lack of knowledge concerning the enzymes involved in polyP metabolism (Desfougères et al., 2020). Many studies have relied on the addition of polyP to cell culture media in order to stimulate signaling from extracellular receptors. Here we demonstrate for the first time that internal accumulation of polyP results in changes at both the transcriptome and proteome levels. Proteomic changes include alterations in protein abundance, but also activation of conserved signaling cascades and mislocalization of a subset of nuclear proteins **(Fig. 7)**. Our work provides a novel resource for studying the impact of mammalian polyP in more depth.

**Figure 7.**
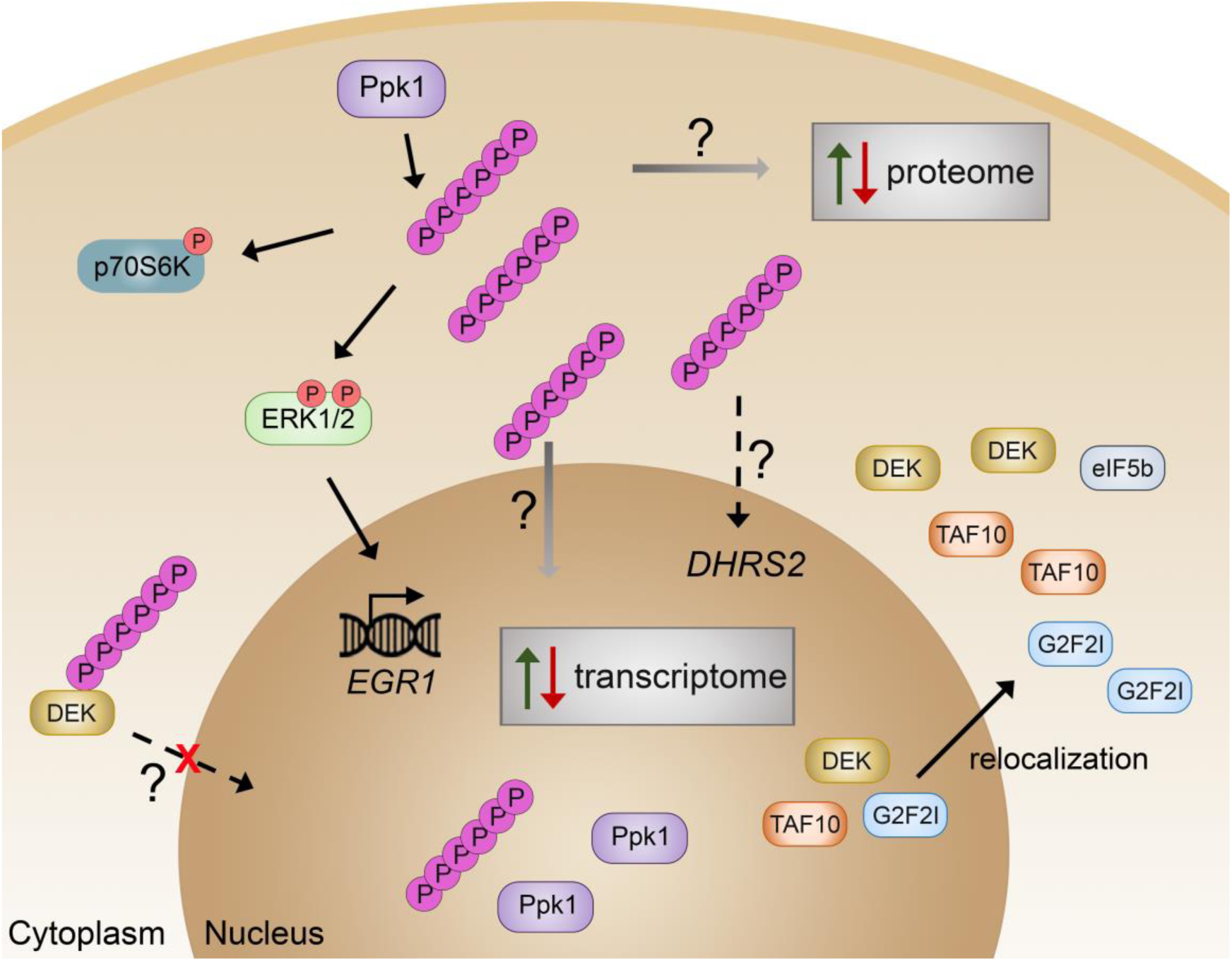
Model. PolyP produced via Ppk1 accumulates inside the cell and results in changes to the transcriptome and the proteome, and relocation of several nuclear proteins. The question marks represent mechanisms still to be explored.

We found that ectopic expression of *ppk1+* results in the production of polyP throughout the cell. Interestingly, the pattern of localization of polyP does not mirror the localization of HA-Ppk1. For example, while most HA-Ppk1 localized to the nucleus/cytoskeleton, this compartment actually contained the lowest amount of polyP. This may reflect either a reduced ability to make polyP or variations in polyphosphatase activity within the nucleus relative to other cellular compartments. It is also possible that polyP synthesized in one cellular compartment could be transported to other organelles. The transport of polyP within cells is poorly understood and represents an important area for future research.

PolyP chains synthesized by Ppk1 in mammalian cells are of the long chain type (>130 phosphate moieties), with the longest chains found in the nuclear/cytoskeleton compartment. These are similar to the polyP synthesized by *E. coli* under stress conditions and qualitatively different from the more heterogeneous chains synthesized by the vacuolar bound VTC complex in budding yeast. A wide range of sizes of polyP chains have been reported in mammalian tissues and cells, ranging from 2 - >5000 Ps in length (Gabel and Thomas, 1971; Kumble and Kornberg, 1995; Pisoni and Lindley, 1992). Short chains (60 – 100 Ps) have been reported in human platelets (Müller et al., 2009; Ruiz et al., 2004). The longest chains have been reported in specific cell and tissue types, such as the lysosomes of human fibroblasts (100-600 Ps) (Pisoni and Lindley, 1992), the rodent brain, and in cell lines such as HEK293s where chains were found to predominantly be 800 residues long (Kumble and Kornberg, 1995). This suggests that the chains made via expression of Ppk1 are reflective of the sizes found in some mammalian cell types.

Notably, the amounts of polyP synthesized in cells transfected with ectopic *ppk1+* are much greater than polyP levels from untransfected cells. While this remains an important caveat of our study, it is possible that untransfected cells contain localized pools of highly concentrated polyP that could impact pathways similar to those uncovered here. The Ppk1 system also provides a method to model the impact of high concentrations of intracellular polyP delivered via nanoparticles, which is being investigated as a method to deliver polyP intracellularly (Fernandes-Cunha et al., 2018). Long-chain polyP produced by Ppk1 may mimic the impact of bacterial long-chain polyP that could be released during infection and taken up by host cells.

We investigated the impact of polyP accumulation on both the human transcriptome and proteome with RNA-seq and mass spectrometry **(Fig. 7)**. This work is the first example of wide-spread changes to the cellular landscape via increased internal polyP synthesis. A previous study examined the impact of external polyP application to SaOS-2 cells using microarrays and found 23 upregulated and 11 downregulated genes that showed expression changes (p-value = 0.05, F.C≤1.5) after a 3 hour incubation with polyP (65 Ps) (Lui et al., 2016). The genes from this study with greatest expression changes (*IL11* and *SLC30A1*) did not show a significant change with Ppk1 expression. However, in agreement with our work, the authors did find significant upregulation of *EGR1* mRNA and an increase in phosphorylation of ERK1/2. Finally, several genes (*ID3*, *ID1*, *MGAM*), were upregulated in the microarray experiment but downregulated in our RNA-seq. This suggests that the mode of exposure (internal vs. external) of polyP is a critical factor in the control of polyP-regulated pathways. Other experimental factors between the two studies could also contribute to the observed differences such as the length of polyP (65 Ps vs. 130 Ps), or exposure time to polyP (3 vs. 48 hours).

Our mass spectrometry analysis revealed that internal polyP accumulation resulted in 14 proteins with significant changes in levels as compared to the controls. Moreover, we uncovered >100 proteins, termed singletons, that showed changes in their levels with polyP, but could not be included in our statistical analysis because they were only detected in 3 or 4 of our biological replicates, instead of all 5 in both conditions. One such example is DHRS2, which showed drastic changes in mRNA levels and was completely undetected in all five Ppk1 biological replicates. DHRS2 protein downregulation via shRNA was previously shown to decrease p53 phosphorylation at Ser15 (Zhou et al., 2018). We observed no significant change in phosphorylated p53 (Ser15) levels accompanying polyP-induced DHRS2 downregulation. This difference could be due to several reasons. First, our study was carried out in HEK293T cells, whereas the silencing of DHRS2 and decrease of p53 (Ser15) phosphorylation was in KYSE510 and KYSE30 cells, both from esophageal squamous cell carcinoma. Additionally, it is possible that signaling cascades impacted by polyP do not reflect “canonical” pathways. It is intriguing to consider that polyP’s effect on some pathways, such as those regulated through DHRS2, may only be apparent under conditions such as stress or infection.

Of our top candidates (FDR<15% and singletons detected in 5 out of 5 replicates in one condition), 3 proteins (RPPB1, MCM3 and DHX9) have PASK motifs, which may be hotspots for lysine polyphosphorylation (Bentley-DeSousa et al., 2018). It is intriguing to speculate that stability of these proteins may be altered by polyphosphorylation. PolyP may also act non-covalently to modulate stability via protein folding (Xie and Jakob, 2019). Finally, polyP may impact protein abundance indirectly by regulating translation or turnover via the ubiquitin proteasome.

Previous work found that external application of polyP activates the ERK1/2 signaling pathway and TOR signaling pathways by interacting with the RAGE and P2Y1 receptors on the exterior of the cell (Hassanian et al., 2015). Here we show that these same pathways can be activated by internal production of polyP **(Fig. 7)**. While we cannot completely exclude the possibility that polyP is being exported to interact with external receptors, we favor a model where polyP functions by promoting activation of upstream regulators. For example, Wang *et al,* previously showed that polyP directly interacts with mTOR to increase its activity (Wang et al., 2003). PolyP may have a similar impact on the RAS, RAF and MEK kinases that sit upstream of the ERK1/2-EGR1 signaling cascade investigated here. Indeed, previous work by the Saiardi lab identified H-RAS as a candidate polyP interacting protein (Azevedo et al., 2018). It is also possible that polyP impacts this pathway indirectly. Given our findings with the ERK1/2-EGR1 pathway, it is perhaps surprising that we did not observe any major changes in cell cycle, autophagy, or apoptosis markers. An effect on these processes may become apparent under specific conditions.

We found that polyP accumulation coincided with the relocalization of several known chromatin bound proteins, GTF2I (Makeyev et al., 2012), DEK (Alexiadis et al., 2000; Waldmann et al., 2002), and TAF10 (Soutoglou et al., 2005) from the nuclear/cytoskeleton fraction. This group includes factors previously described as targets of lysine polyphosphorylation (eIF5b, GTF2I and DEK) **(Fig. 7)**. Interestingly, Ppk1 expression resulted in a decrease in nuclear/cytoskeleton eIF5b without a corresponding increase in other fractions. PolyP may promote the degradation of eIF5b in this fraction. For DEK, we observed a corresponding increase in protein accumulation in the cytoplasm and the mitochondrial/organelle fractions. Since polyP produced by Ppk1 is sufficient to cause polyphosphorylation of DEK in all fractions, it is tempting to speculate that polyphosphorylation might regulate DEK localization directly, perhaps by preventing nuclear import. Indeed, DEK’s predicted nuclear localization signal is immediately adjacent to one of its three poly-acidic serine and lysine rich (PASK) regions that are common sites for polyphosphorylation (Bentley-DeSousa et al., 2018). Our work provides a unique resource to explore polyphosphorylation and other aspects of polyP biology in greater detail.

## Materials and Methods

### Cell lines & maintenance

The HEK293T cell lines used in this study were obtained from the American Type Culture Collection (ATCC CRL-3216). Cells were grown at 37 °C and 5% CO2 to ∼70% confluency in DMEM (Wisent, 319-015-CL) supplemented with 10% Fetal Bovine Serum (Wisent, 080-150) and 1% Penicillin-Streptomycin-Amphotericin B solution (Wisent, 450-201-EL). For the RNA-seq, ERK1/2, and p70SK6 experiments cells were grown with the addition of 1 mM sodium pyruvate (Sigma, S8636) in the media. All cell lines used were regularly tested for mycoplasma contamination using a PCR Mycoplasma Detection Kit (Applied Biological Materials, G238).

### Yeast and bacterial strains

Wild-type and *vtc4*Δ::NatMX *S. cerevisiae* strains containing empty vectors allowing growth in SC-Uracil minimal media are of the BY4741 S288C background with the genotype *MAT*a *leu2Δ0 ura3Δ0 met15Δ0 his3-1*. Strains were grown in SC-URA media at 30 °C to maintain the plasmids. For bacterial polyP extraction, WT MG1655 *E. coli* of genotype *F-lambda-ilvG-rfb-50 rph-1* was grown as described below for ‘polyphosphate analysis’.

### Plasmids

Yeast strains contain the plasmid p416 (ATCC) to allow growth in synthetic media lacking uracil. Mammalian *ppk1+* expression vector contains the *E. coli ppk1+* sequence cloned into a version of pcDNA3.1(-) containing a bleomycin resistance marker. This plasmid has been described previously (Bentley-DeSousa et al., 2018) and is available from Addgene (Plasmid #108850). Mammalian HA*ppk1+* expression vector contains the *E. coli ppk1+* sequence including a HA-tag on the N-terminal, cloned into a version of pcDNA3.1(-) containing a bleomycin resistance marker. This plasmid will be available on Addgene.

### Polyphosphate analysis

The polyP isolation protocol was adapted from (Bru et al., 2017). HEK293T cell pellets were resuspended in 400 µL of LETS buffer (100 mM LiCl, 10 mM EDTA, 10 mM Tris-HCl pH 7.4, 0.2% SDS) kept at 4 °C. Samples were mixed with 600 µL of phenol pH=8 and 150 µL of water in a screw cap tube, and vortexed for 20 seconds. They were then heated for 5 minutes at 65 °C then cooled down on ice. 600 µL of chloroform were added followed by 20 seconds of vortexing. Samples were then centrifuged at room-temperature for 2 minutes at 13 000 x g and the top layer/water phase was transferred to a new screw cap tube containing 600 µL of chloroform. Samples were vortexed and centrifuged again for 2 minutes at 13 000 x g, and the top layer was transferred to a clean 1.5 mL tube. 2 µL of 10 mg/mL RNAse A and 2 µL of 10 mg/mL DNAse I were added followed by 1 hour incubation at 37 °C. Samples were transferred to pre-cold 1.5 mL tubes containing 1 mL of 100% ethanol and 40 µL of 3 M sodium acetate pH=5.3. polyP was left to precipitate at −20 °C for at least 3 hours. Samples were centrifuged for 20 min at 13 000 x g at 4 °C and the supernatant was discarded. PolyP pellets were washed with 500 µL of 70% ethanol and centrifuged for an additional 5 minutes. The supernatant was discarded and the pellets air dried for a few minutes before being resuspended in 20 µL of water.

Yeast polyP preps were from cells diluted at OD600 = 0.2 in YPD and grown at 30 °C until the OD reached 0.8-1. The equivalent of 6 OD600 units were used in the prep and the final polyP pellet was dissolved in 30 µL of water. For *E. coli* polyP preps, overnight cultures were diluted to OD600 = 0.1 in 20 mL of LB media in the absence of antibiotics and grown at 37 °C. Cells were grown to exponential phase (OD600 = 0.4-06) prior to harvesting (control) or switching to MOPs minimal media to induce starvation. OD600 units were harvested for control samples and these cells were washed with cold water, pelleted and frozen on dry ice. For starvation samples, 6 OD equivalent of cells were washed once with 1 X PBS [100 mL of 10 × 1L Stock (54.36 g NaH_2_PO_4_ (Sigma-Aldrich S9638-1KG), 160 g NaCl (Fisher Scientific, AM9759), 4 g KCl (Sigma-Aldrich 746436-500G), 9.6 g KH_2_PO_4_ (Sigma-Aldrich P0662-500G), fill to 1 L ddH2O), and 900 mL ddH2O], resuspended in 10 mL of 1 X MOPS (Sigma-Aldrich M1254-1KG) media and incubated at 37 °C for and additional 2 hours. Starved cells were harvested similar to control cells by centrifugation, washing with cold water, and freezing on dry ice. Cells were then processed as described for HEK293T cells.

Extracted polyP with loading dye (10 mM Tris-HCl pH 7, 1 mM EDTA, 30% glycerol, bromophenol blue) was run on a 15.8% acrylamide TBE-Urea gel (5.8 M Urea - 5.25 g Urea, 7.9 mL 30% acrylamide, 3 mL 5 X TBE, 150 µL 10% APS, 15 µL TEMED) at 100 V for 1 hour and 45 minutes in 1 X TBE buffer.

Negative DAPI staining protocol was adapted from (Smith and Morrissey, 2007). Briefly, gel was incubated for 30 minutes in fixative solution (25% methanol, 5% glycerol, 50 mM Tris base - pH 10.5) containing 2 μg/mL of DAPI at room temperature. Following this incubation, the gel was destained for 1 hour in fixative solution with a replacement of fresh fixative solution after 30 minutes. Gel was exposed on a 365 nm UV transilluminator to photobleach DAPI bound to PolyP. This step required adjustable times (average several minutes) based on the amount of polyP. Finally, the gel was imaged using a Chemi Doc at 312 nm.

### Transfection and lysate harvest

HEK293T cells were grown to ∼70% confluency and transiently transfected with 2-28 μg of either control plasmid (pcDNA3.1-) or *ppk1+* expression plasmids (pcDNA3.1-*ppk1+*), (pcDNA3.1-HA*ppk1+*) using Lipofectamine 2000 (Thermo Fisher Scientific, 11668019) or Lipofectamine LTX Reagent (Thermo Fisher Scientific, 15338100). Transfected cells were grown for 48 hours in OptiMeM (Fisher Scientific, 31985-070) and then harvested.

#### Protein

To isolate total protein, cells were washed with ice-cold 1 X PBS, scraped, and lysed in ∼200 μL (3.5 cm plate) or ∼1 mL (10 cm plate) of ice-cold radioimmunoprecipitation assay buffer (RIPA) [50 mL ice cold lysis buffer [10 mM Tris-HCl pH 7.4, Hydrochloric acid (Fisher Scientific, A144 212), 100 mM NaCl (Fisher Scientific, AM9759), 1 mM EDTA (Sigma-Aldrich, 03690-100ml), 1% IGEPAL (Sigma-Aldrich, I8896-50ML), 0.5% Sodium Deoxycholate (Sigma-Aldrich, D6750-25G), 0.1% SDS (Thermo Fisher Scientific, 15525-017)], 500 μL Sodium Fluoride (0.5 M, Sigma-Aldrich, 201154-5G), 500 μL Glycerol-2-Phosphate (1 M), 500 μL Nicotinamide (1 M Sigma-Aldrich, N3376-100G), 500 μL PMSF (1 mg/mL Sigma-Aldrich, P7626-1G), 500 μL Sodium Butyrate (1 M Sigma-Aldrich, 303410-100G), and 1 cOmplete protease inhibitor tablet (Roche, 04693159001)]. Following a 15 minute incubation on ice, the cells were centrifuged for 15 minutes at 13,500 RPM. Supernatant was collected and either used immediately for analysis with immunoblotting or stored at −20°C.

To isolate the cell population for cell fractionation experiments, cells were washed with ice-cold 1 X PBS, scraped, and centrifuged for 5 minutes at 4°C at 350 RCF. Cell pellet was then resuspended in 1 X PBS and cell fractionation was carried out using the Cell Fractionation Kit (Cell Signaling Technology, 9038) according to manufacturer’s protocol.

#### RNA

To isolate total RNA, cells were washed with ice-cold 1 X PBS and scraped in 1 mL of ice-cold TRIzol (Thermo Fisher Scientific, 15596026). Cells were then briefly vortexed and incubated at room temperature for 5 minutes. Following incubation, 200 µL of chloroform (Sigma-Aldrich, 472476-500ml) was added and cells were vortexed for 30 seconds and incubated for 3 minutes at room temperature. Cells were then centrifuged for 15 minutes at 4°C at 12,000 RCF. The aqueous phase was transferred to a new tube and 500 μL of ice-cold isopropanol (Sigma-Aldrich, 278475-1L) and 2 μL of Glycoblue (Thermo Fisher Scientific, AM9515) was added. Mixture was incubated for 10 minutes at room temperature and then centrifuged for 15 minutes at 4°C at 12,000 RCF. Isopropanol was carefully removed and pellet was resuspended in 500 μL of ice-cold ethanol (Commercial Alcohols, P016EAAN). Following a 10 minute centrifugation at 4°C at 7, 600 RCF the pellet was air-dried for 5 minutes at room temperature. Dried pellet was resuspended in 25 μL of RNase-free water. All RNA work was carried out with filter pipette tips.

### Western blotting

Following protein harvest and preparation (cell fractionation) samples were analyzed by western blotting using SDS-PAGE and NuPAGE. 5 X Laemmli Sample Buffer was added to the protein samples prior to boiling at 95°C for 2 minutes and briefly centrifuged. Fraction samples used for cell fractionation experiments were boiled for 5 minutes and centrifuged for 3 minutes at 15,000 x RCF, as recommended according to manufacturer’s protocol. All antibodies and dilutions used in this study are described in **Supplemental Table 1**. Please note that SDS-PAGE gels do not resolve polyP-induced protein shifts (Azevedo et al., 2015; Bentley-DeSousa et al., 2018).

#### SDS-PAGE

Samples were separated using 10-15% SDS-PAGE gels or 4–20% Criterion TGX Stain-Free Protein Gel (Bio-Rad, 5678095). Gels were run at 100 V using 1 X SDS-PAGE Running Buffer [100 mL of 10 × 1 L Stock [30.2 g Tris Base (Fisher Scientific, BP152-5), 188 g Glycine (Sigma-Aldrich, G7126-500G), 10 g SDS (Thermo Fisher Scientific, 15525-017)] and 900 mL ddH2O] and transferred to Polyvinylidene fluoride membranes (Bio-Rad, 162-0177) at 85 V for 1 hour in 1 X SDS-PAGE Transfer Buffer [100 mL of 10 × 1L Stock [30.275 g Tris Base (Fisher Scientific, BP152-5), 166.175 Glycine (Sigma-Aldrich, G7126-500G), 200 mL Methanol (Fisher Scientific, A412P-4)] and 700 mL ddH2O.

#### NuPAGE

Samples were separated using 4-12% Bis-Tris NuPAGE gels (Thermo Fisher Scientific, NP0336BOX) at 200 V using 1 X NuPAGE Running Buffer [50 mL of 20 × 1 L Stock [209.2 g MOPS (Sigma-Aldrich, M1254-1KG), 121.1 g Bis-Tris (Sigma-Aldrich, B9754-1KG), 20 g SDS (Thermo Fisher Scientific, 15525-017), 12 g EDTA (Sigma-Aldrich, 03690-100ml)] and 950 mL ddH2O]. The gels were then transferred to Polyvinylidene fluoride membranes at 85V for 1 hour using 1 X NuPAGE Transfer Buffer [50 mL of 20 × 1 L Stock [81.6 g Bicine (Sigma-Aldrich, B3876-250G), 104.8 g Bis-Tris (Sigma-Aldrich, B9754-1KG), 6 gEDTA (Sigma-Aldrich, 03690-100ml)], 200 mL Methanol (Fisher Scientific, A412P-4) and 750 mL ddH2O].

#### Immunoblotting

In general, membranes were blocked for 30 minutes with either 5% milk or BSA (VWR, 97061-416) diluted in 1 X TBST [100 mL of 10 × 1L Stock (90 g Tris Base (Fisher Scientific, BP152-5), 88 g NaCl (Fisher Scientific, AM9759), 2 g KCl (Sigma-Aldrich 746436-500G), adjust pH with HCl to 7.5, fill to 1 L ddH2O), 10 mL of 10% Tween 20 (Fisher Scientific BP337500) and 850 mL ddH2O] under gentle agitation at room temperature. Primary antibodies were diluted either with 5% milk or BSA diluted in 1 X TBST and incubated with membrane at 4°C overnight or 1 hour at room temperature. Following use, primary antibodies were stored at 4°C with 1% sodium azide. After primary antibody incubation, membranes were washed with 1 X TBST for 10 minutes at room temperature 3 times followed by a 30 minute incubation with horseradish peroxidase-conjugated secondary antibodies. Following 3 more 10 minute washes with 1 X TBST at room temperature, membranes were incubated with Chemiluminescence HRP Substrate, Luminata Forte (Fisher Scientific, WBLUF0500) and exposed to autoradiography film (Harvard Apparatus Canada, DV-E3018). Of note, Vimentin blots were incubated with fluorescent IR Dye 800CW secondary antibodies for 1 hour, washed in 1 X TBST 3 times (10 minutes), and visualized using a LI-COR Odyssey imaging system.

#### Protein screen

Proteins that were screened (11 proteins) for fractionation experiments were Hsp90, Nucleolin, CDC5L, Cyclin A, Cyclin B1, H3, KAT2A/Gcn5, Lamin A/C, p53, RALY, TAF10. All antibodies and dilutions are listed in **Supplemental Table 1**.

### RT-qPCR setup and analysis

cDNA synthesis was carried out with isolated total RNA using 5 X All-In-One RT MasterMix kit (Applied Biological Materials, G490) according to manufacturer’s protocol. Reverse transcription conditions: 25°C for 10 minutes, 42°C for 50 minutes, and 85°C for 5 minutes. Newly synthesized cDNA was used immediately or stored at −20 °C. Primer design for qPCR primers was completed using NCBI Primer-BLAST web-based program with standard parameters (product size between 80-200 bps, T_m_=∼60°C (±3°C), GC content ≤50%). qPCR reactions were carried out using SYBR Green Supermix (Bio-Rad, 1708880) according to manufacturer’s protocol with the following conditions: 95°C 2 minutes, (95°C 30 sec, 60°C 30 sec, 72°C 45 sec)×40 cycles, 72°C 10 minutes). Standard curves were completed for every qPCR experiment and to confirm accurate primer pair efficiency for each primer pair used in this study **(Supplemental Table 3)**. All qPCR reactions were completed in 3 technical replicates and included ≤3 biological replicates and normalized to a control gene. The changes in mRNA expression is represented as fold change and was calculated using the ΔΔCt method (Schmittgen and Livak, 2008).

### RNA-sequencing (RNA-seq)

#### Preparation of RNA-seq samples

RNA-seq was carried out on HEK293T cells transiently transfected in either CTRL or Ppk1-expressing conditions. Three biological replicates were used for this experiment. Following a 48 hour transfection, RNA for RNA-seq was extracted using an miRNeasy Mini Kit (Qiagen, 217004). Protein was also isolated from each sample (detailed in “Transfection and lysate harvest”) and analyzed on NuPAGE gels to confirm detection of shifts on target proteins DEK and MESD **(Fig. 2)**. RNA pellets were stored at −80 °C and shipped on dry ice to Genome Quebec Innovation Centre for RNA sequencing.

The RNA-seq was carried out at Genome Quebec as follows. Total RNA was quantified with a NanoDrop Spectrophotometer ND-1000 (NanoDrop Technologies, Inc.). RNA integrity was assessed on a 2100 Bioanalyzer (Agilent Technologies). To generate libraries from 250 ng of total RNA, the following steps were taken. First, mRNA enrichment was performed using the NEBNext Poly(A) Magnetic Isolation Module (New England BioLabs followed by cDNA synthesis using the NEBNext RNA First Strand Synthesis and NEBNext Ultra Directional RNA Second Strand Synthesis Modules (New England BioLabs). Next, library preparation was completed using the NEBNext Ultra II DNA Library Prep Kit for Illumina (New England BioLabs). The adapters and PCR primers used were purchased from New England BioLabs. Quantification of libraries was carried out using the Quant-iT PicoGreen dsDNA Assay Kit (Life Technologies) and the Kapa Illumina GA with Revised Primers-SYBR Fast Universal kit (Kapa Biosystems) with the average size fragment determined with a LabChip GX (PerkinElmer) instrument. Libraries were first normalized, then denatured in 0.05 N NaOH followed by dilution to 200 pM and finally neutralized using HT1 buffer. Clustering was carried out on an Illumina cBot. The flowcell was run on a HiSeq 4000 for 2×100 cycles in paired-end mode following the manufacturer’s instructions. Used as a control was a phiX library mixed with libraries at 1% level. The Illumina control software used was HCS HD 3.4.0.38 and RTA v. 2.7.7 was the program used for real-time analysis. Finally, the bcl2fastq v2.20 program was used to demultiplex samples and generate fastq reads.

Analysis of RNA-seq was completed by Canadian Center for Computational Genomics (C3G) as follows. Paired-ends sequencing reads were first clipped for adapter sequence and then trimmed for minimum quality (Q30) in 3’. Following this, sequencing reads were filtered for minimum length of 32 bp using Trimmomatic (Bolger et al., 2014). The surviving read pairs were then aligned to the Genome Reference Consortium Human Build 38 using STAR (Dobin et al., 2013). This alignment used the two-passes method. Gene quantification (gene-level count-based) against Ensembl annotations was carried out using HT-seq count (Anders et al., 2015) (mode used: intersection-nonempty). Following gene quantification, exploratory analysis was then conducted with various functions and packages from R and the Bioconductor project (Huber et al., 2015). To complete differential expression both edgeR (Robinson et al., 2010) and DESeq (Anders and Huber, 2010) were used. Significance was determined by DESeq adjusted p-values < 0.05. Gene Ontology terms were tested for enrichment with the GOseq (Young et al., 2010) R package (FDR threshold of 0.05). As expected, *Ecppk1* sequences were detected in Ppk1 expressing samples, but not in the CTRL samples. Each processing step described above was accomplished through the GenPipes framework (Bourgey et al., 2019).

### Liquid Chromatography – Mass Spectrometry

#### Preparation of mass spectrometry samples

Mass spectrometry was carried out on HEK293T cells transiently transfected in either CTRL or *ppk1+* plasmid expressing conditions (five biological replicates for each condition). Following a 48 hour transfection (detailed in “Transfection and lysate harvest”), proteins were harvested, pelleted, and frozen at −80 °C. A small amount of protein was used immediately for analysis on NuPAGE gels to confirm detection of shifts on target proteins DEK and MESD. Protein pellets were shipped overnight on dry ice to the UC Davis Genome Center Proteomics Core Facility. Extracts were prepared via sonication in 5% SDS, 50mM TEAB, supplemented with protease and phosphatase inhibitor tablets (Roche). Extracts were clarified via centrifugation at 15,000xg for 10min @ 4 °C prior to trypsin digestion using S-Trap Mini spin columns (PROTIFI) as per manufacturer’s instructions.

#### Liquid chromatography and mass spectrometry analysis

LC separation was carried out on a Proxeon Easy1200 HPLC (Thermo Fisher Scientific) with a Proxeon nanospray source (operating in positive ionization mode). After LC separation, digested peptides were then reconstituted in 2% acetonitrile/0.1% trifluoroacetic acid. 1 μg of each sample was then loaded onto a 100 μm×25 mm Magic C18, 100 Å 5 μm reverse phase trap wherein samples were desalted online, and directly eluted onto an analytical column (maintained at 40⁰ C) for on-line separation. The analytical column used was a 75μm×150 mmMagic C18 200 Å 3μm reverse phase column. Peptides were then eluted into the mass spectrometer using a 2-buffer gradient (Buffer A: 0.1% formic acid in water and Buffer B: 0.1% formic acid in 100% acetonitrile) at a flow rate of 300 nL/min.

A 144 minute gradient was run as follows: 2 to 25% Buffer B (90 minutes), 25% to 40% Buffer B (26 minutes), 40 to 100% Buffer B (4 minutes), held at 100% Buffer B (3 minutes). Finally, ending with 2% Buffer B (21 minutes). The analytical column used tapers off into the electrospray emitter. The peptides were then ionized (spray voltage 2.0 kV). An Orbitrap Q Exactive Plus mass spectrometer (Thermo Fisher Scientific) was used to collect the mass spectra. Mass spectra was collected in a data-dependent mode with the following: MS survey scan (m/z range 350–1,600), 70,000 resolution, target of 1 × 106 ions or a maximum injection time of 30 ms, 15 MS/MS scans (where the top 15 ions in the MS spectra were subjected to high-energy collisional dissociation, and normalised fragmentation energy of 27%. The MS/MS spectra were obtained at a resolution of 17,500 and a target of 5 × 104 ions or a maximum injection time of 50 ms. The isolation mass window used for precursor ion selection was 1.6 m/z, with accepted charge states 2-4, and a normalized collision energy of 27% used for fragmentation. Finally, precursor ion dynamic exclusion was set to 20 seconds.

#### Protein identification

LC-MS/MS RAW data files were converted to mzML files using MSConvert from ProteoWizard (Adusumilli and Mallick, 2017). mzML files were then searched using X! Tandem (Bjornson et al., 2008) against the UniProt human protein sequence database (downloaded March 4, 2019, from uniprot.org;) (Consortium, 2015) using a target-decoy strategy. The database contained 73,948 protein sequences amended with 49 potential contaminants from the cRAP database of common laboratory contaminants (www.thegpm.org/crap) and an equal number of reversed decoy sequences. The database search was performed using trypsin as the digestive enzyme and allowing for one missed cleavage, as well as carbamidomethylation of Cys as a fixed modification. Up to five variable modifications were allowed per peptide, including oxidation of Met and Trp, as well as n-terminal acetylation. Instrument parameters and match tolerances were set to Orbitrap defaults. A protein identification required at least one non-redundant peptide (unique) identification. Protein identifications were grouped and clustered using Scaffold (v4.1, Proteome Software). Protein and peptide identification confidence threshold was set to 1% FDR and were matched between runs and second peptides enabled.

#### Protein differential expression analysis

Protein and protein cluster spectral counts from Scaffold were normalized based on the total number of spectral counts in each replicate. Protein differential expression was statistically assessed using a two-tailed, two-sample Student’s *t*-test assuming unequal variance, which was performed on normalized spectral counts of proteins or protein clusters detected in all 5 replicates in each condition. This criterium was selected to maximize the accuracy of the *t*-test statistical assessment. An FDR was estimated at different *p*-value thresholds **(Supplementary Table 4)**. On the other hand, singleton proteins are defined as proteins that were identified in at least 3 replicates in one condition and 0 in another (All or None (5 replicates/5 replicates vs. 0 replicates/5 replicates) – Mostly All or None (3 or 4)/5 vs. 0/5) or proteins that were identified in all 5 replicates in one condition but in 2 or less in the other condition (All or Mostly None (5/5 vs. (1 or 2)/5) **(Supplementary Table 4)**.

#### Gene Ontology enrichment analysis

Differentially expressed proteins (FDR <15%) and singleton proteins were investigated for Gene Ontology enrichments using Ontologizer (Bauer et al., 2008). All quantified proteins were used as background.

#### Protein-protein interaction network construction with STRING

The set of differentially expressed proteins (FDR <15%) and singleton proteins was used to query their protein-protein interactions from the STRING database (Szklarczyk et al., 2019). Protein-protein interactions derived from experiments and databases with a medium confidence level were used to build the network. All proteins without any protein-protein interactions with a protein in the query set were not included in the network. Gene Ontology term overlay on the network was performed using STRING.

### Data availability

RNA-seq data has been deposited into the GEO database under the identifier GSE149011.

Mass spectrometry data have been deposited to the ProteomeXchange Consortium (Deutsch et al., 2016) via the PRIDE partner repository (Perez-Riverol et al., 2019) with the dataset identifiers PXD018896 and 10.6019/PXD018896. Reviewers can access the data on PRIDE using the following credentials: username: and password: GcJJmmdb.

## Acknowledgments

We thank the Downey lab members for critical reading of the manuscript. We thank T. Shiba (Regentiss, Japan) for the generous gift of polyP standards. This study was funded by a Canadian Institutes of Health Research Project Grant to MD (PJT-148722). E. B.-C. was supported in part by a postdoctoral fellowship from The Natural Sciences and Engineering Research Council of Canada (NSERC). I.A. was supported in part by a NSERC Undergraduate Student Research Award. M.L.A. holds a NSERC Discovery Grant.

The Canadian Center for Computational Genomics (C3G) is a Genomics Technology Platform (GTP) supported by the Canadian Government through Genome Canada.

---

**Supplementary Figure 1.**
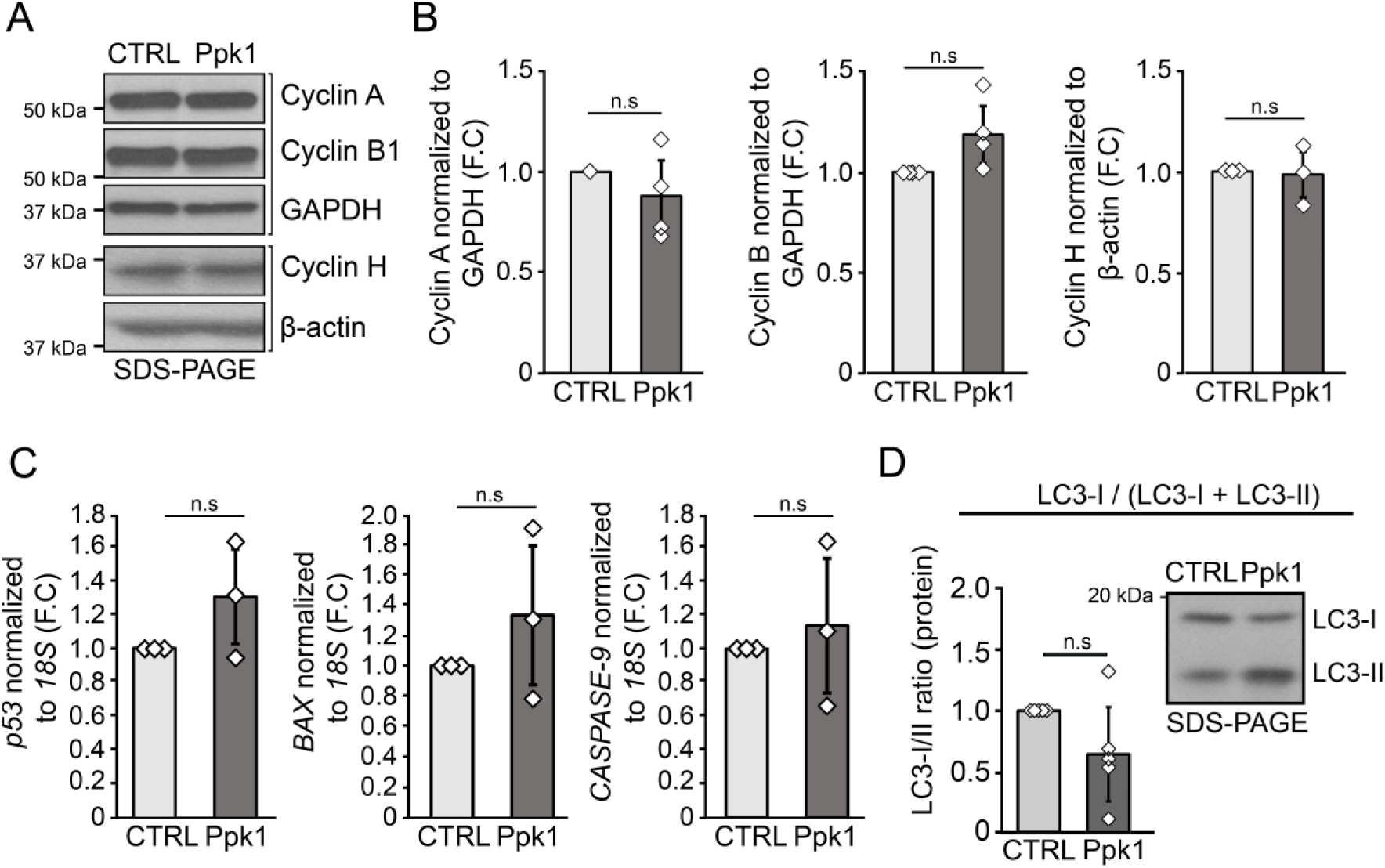
No significant impact of polyP on cell cycle, apoptosis, or autophagy. RNA and protein were harvested from HEK293T cells following a 48-hour transient transfection of either a control or *ppk1+* expression plasmid. *(A)* Western blotting analysis by SDS-PAGE in control and Ppk1 conditions cells using antibodies against Cyclin A, Cyclin B1, GAPDH, Cyclin H, and β-actin. Image is representative of N≤3. Brackets indicate the separation of biological replicates shown. For each bracketed image, the same biological sample was used from different blots analyzed in parallel. GAPDH or β-actin were used as loading controls. *(B)* Semi-quantitative analysis of protein levels shown in panel A. *(C)* RT-qPCR of *p53, BAX, and CASPASE-9* mRNA levels in control and Ppk1 conditions. Changes in mRNA levels are represented by fold change (F.C). *(D)* Western blotting analysis by SDS-PAGE and semi-quantitative analysis of protein levels of LC3 in control and Ppk1 conditions cells. P-values are shown as numerical values or if p≤0.05 as non-significant (n.s). Statistical tests performed were one-sample t-Test (unequal variances) where error bars represent standard deviation. Image shows results from one biological replicate which is representative of N≤3.

**Supplementary Figure 2.**
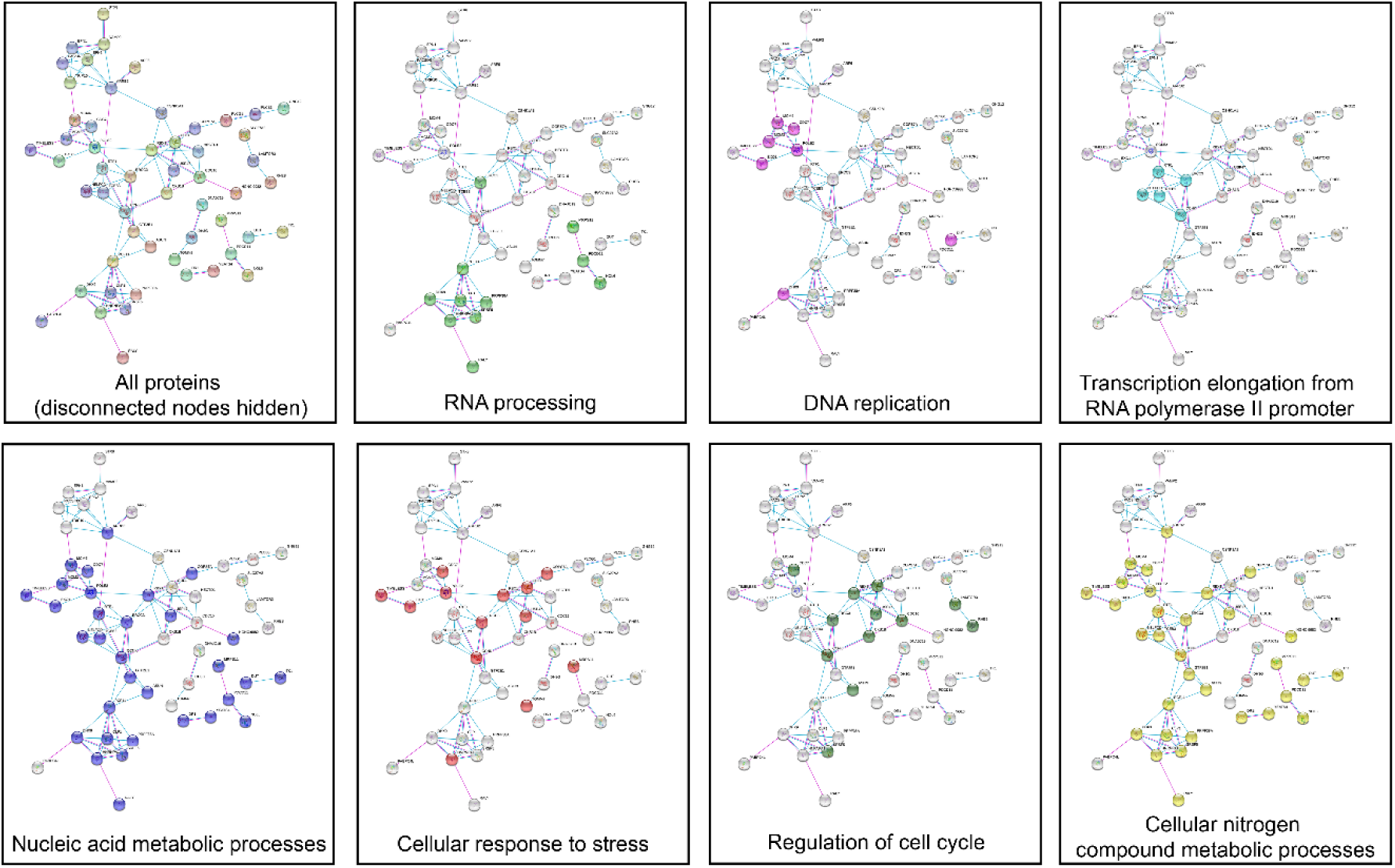
STRING. Proteins classified as significantly differentially expressed and singletons based on the mass spectrometry experiment **(Supplemental Table 4 and materials and methods)** were analyzed using the web-based STRING program v11.0 (Szklarczyk et al., 2019). Network was generated using known interactions (curated databases and experimentally determined) with disconnected nodes hidden from network. GO-terms associated with proteins shown here belong to the “biological processes” sub-ontology. Nodes in color with the terms ‘RNA processing’, ‘DNA replication’, and ‘transcription elongation from RNA polymerase II promoter’ were nodes forming isolated clusters.

**Supplementary Table 1.** Antibodies used in this study.

| **Antibody target** | **Dilution** | **Source** | **Identifier** |
| --- | --- | --- | --- |
| Primary antibodies |  |  |  |
| Beta-Actin HRP-conjugate | 1:40,000 | Santa Cruz | Sc-47778 |
| CDC5L | 1:1,000 | BD Transduction Laboratories | 612362 |
| Cyclin A | 1:1,000 | Santa Cruz | SC-596 |
| Cyclin B1 | 1:1,000 | Santa Cruz | SC-752 |
| Cyclin H | 1:1,000 | Santa Cruz | SC-855 |
| DEK | 1:1,000 | Santa Cruz | 136222 |
| DHRS2 | 1:1,000 | Cedarlane | GTX123431-25UL |
| EGR1 | 1:500 | Cell Signalling Technologies | 44D5 |
| eIF5b | 1:200 | Santa Cruz | 393564 |
| ERK1/2 | 1:1,000 | Cell Signaling Technologies | 4695S |
| GAPDH HRP | 1:40,000 | Abcam | Ab85760 |
| H3 | 1:10,000 | Abcam | Ab1791 |
| HA | 1:500 | Santa Cruz Biotechnology | sc-57592 |
| Hsp90 | 1:1,000 | Santa Cruz Biotechnology | Sc-13119 |
| KAT2A/Gcn5 | 1:1,000 | Gift from Dr. Laszlo Tora | 2GC-2C11 |
| Lamin A/C | 1:1,000 | Abcam | Ab108922 |
| LC3 | 1:1,000 | Novus | NB100-2220 |
| MESD | 1:1,000 | Cell Signalling Technologies | 2763 |
| Nucleolin | 1:5,000 | Abcam | Ab22758 |
| p-p70S6K | 1:1,000 | Cell Signaling Technologies | 9234S |
| p53 | 1:1,000 | Cell Signaling Technologies | 2524 |
| p70S6K | 1:1,000 | Cell Signalling Technologies | 2708S |
| p-ERK1/2 | 1:1,000 | Cell Signalling Technologies | 9101S |
| p-p53 (Ser15) | 1:1,000 | Cell Signalling Technologies | 9284 |
| RALY | 1:1,000 | EMD millipore | AB12226E104 |
| TAF10 | 1:1,000 | Gift from Dr. Laszlo Tora | 6TA 2B11 |
| TFII-I | 1:1,000 | Santa Cruz | sc-46670 |
| Vimentin | 1:1,000 | BD Pharmingen | 550513 |
| Secondary antibodies |  |  |  |
| Goat anti-Mouse HRP | 1:10,000 | Biorad | 172-1011 |
| Goat anti-Rabbit HRP | 1:10,000 | Biorad | 170-6515 |
| IR Dye 800CW Goat anti-Mouse IgG | 1:5,000 | LI-COR | 925-32210 |

**Supplementary Table 2.** Downey_RNA-seq_Analysis_Processed_data_2020.

GEO submission identifier: GSE149011

**Supplementary Table 3.**
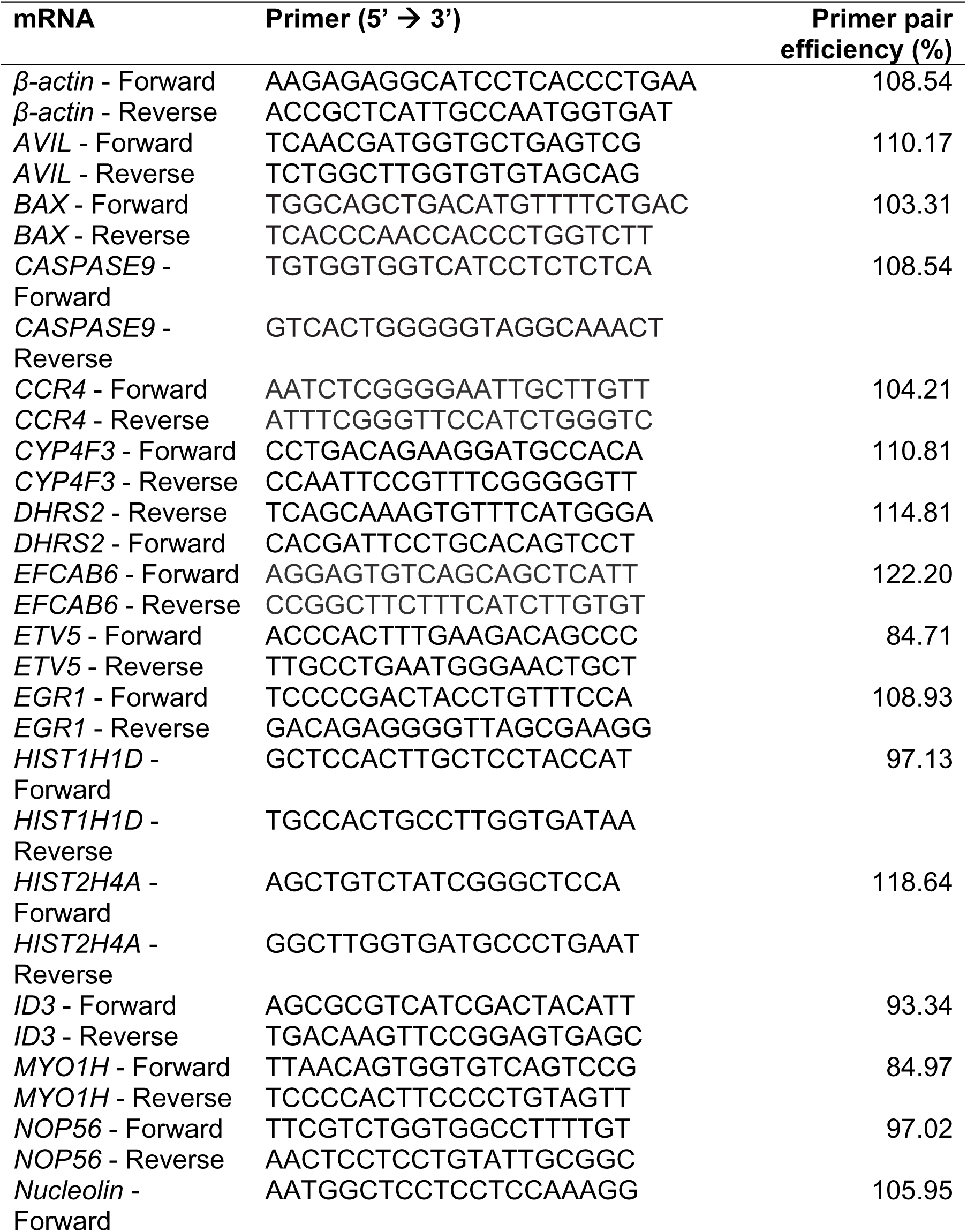
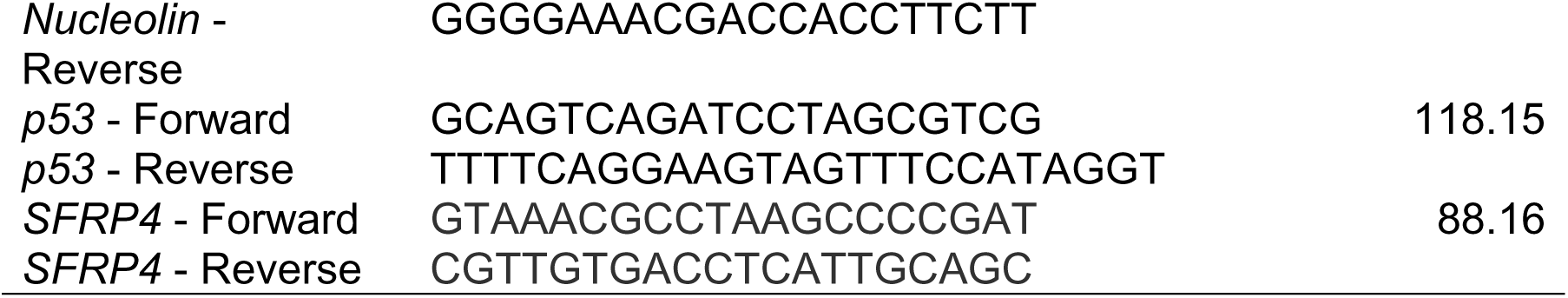
RT-qPCR primers used in this study. Primer design is detailed in materials and methods.

**Supplementary Table 4.** Differential Protein Expression Summary Downey Ppk1 Mass Spectrometry. Normalized spectral counts and statistical analysis of differential expression analyses derived from mass spectrometry data.

PRIDE identifier PXD018896 and 10.6019/PXD018896

## Notes

### Competing Interest Statement

The authors have declared no competing interest.

### Summary of Updates

Simplified manuscript abstract that fits on one page.

